# Intrinsic timescale develops hierarchically from sensorimotor to association cortex in youth and stabilizes in adulthood

**DOI:** 10.64898/2026.04.08.717312

**Authors:** Golia Shafiei, Joëlle Bagautdinova, Valerie J. Sydnor, Dani S. Bassett, Deanna M. Barch, Matthew Cieslak, Yong Fan, Elizabeth Flook, Alexandre R Franco, Gregory Kiar, Audrey C. Luo, Michael P Milham, Linden Parkes, Taylor Salo, Leah H. Somerville, Tien T. Tong, Russell T. Shinohara, Theodore D. Satterthwaite

## Abstract

Intrinsic timescale is a commonly used measure of spontaneous neural dynamics that quantifies the temporal window of processing of neuronal populations. Intrinsic timescale displays a hierarchical cortical organization across multiple species and imaging modalities, with shorter timescales in sensorimotor cortex compared to association cortex. However, less is known about how intrinsic timescale evolves during human brain development and whether its cortical maturation patterns generalize to independent developmental samples. Here we estimate the intrinsic timescale in two independent cross-sectional datasets of youth (HCPD: *n*=565; HBN: *n*=729; age range 8–22 years) and investigate its neurodevelopmental patterns. We find that developmental patterns in the intrinsic timescale follow a hierarchical pattern that recapitulates an axis spanning sensorimotor to association cortices (S–A axis). Our analysis of an independent healthy young adult dataset (HCPYA: *n*=973, age range 22–37 years) underscores the specificity of these developmental findings, suggesting that the intrinsic timescale develops along the S–A axis in youth and stabilizes in adulthood. Together, these results reveal convergence between major axes of cortical organization and development, highlighting intrinsic timescale as a principled marker of hierarchical brain maturation in youth.

## Introduction

Examining the spontaneous neural activity of the human cortex can help reveal how neural populations integrate information over time. Intrinsic timescale is a commonly used measure of spontaneous neural dynamics that captures the temporal window over which neural populations accumulate, maintain, and dissipate information (“memory” of a signal; Hasson et al., 2008; Honey et al., 2012; Murray et al., 2014). Across species, recording techniques, and imaging modalities, intrinsic timescale is hierarchically organized across the cortex in adulthood: sensorimotor regions exhibit short timescales, whereas association cortices show longer integration windows (Murray et al., 2014; Ito et al., 2020; Gao et al., 2020; Raut et al., 2020; Shafiei et al., 2023). This hierarchy aligns closely with major axes of cortical structure, function, and evolution, suggesting shared principles that link local circuit properties to large-scale cortical organization (Margulies et al., 2016; Burt et al., 2018; Huntenburg et al., 2018; Demirtas et al., 2019; Sydnor et al., 2021). Recent developmental findings suggest that the same cortical axes that organize intrinsic timescale in adults, particularly the sensorimotor–association (S–A) axis, also align with brain maturation in youth (Sydnor et al., 2021; Luo et al., 2024; Bero et al., 2026). However, it remains unclear if the development of the intrinsic timescale follows a hierarchical sequence. Here, we evaluate if intrinsic timescale quantified using functional Magnetic Resonance Imaging (fMRI) develops along the cortical hierarchy using multiple large samples of children and adolescents.

Regional variation in intrinsic timescale is thought to arise from underlying cellular and circuit-level mechanisms that shape the temporal integration properties of local neuronal populations (Kiebel et al., 2008; Chaudhuri et al., 2015; Murray et al., 2018; Wang, 2020). At the cellular level, neuronal properties such as intrinsic membrane time constants, ion channel dynamics, and dendritic morphology set a baseline for temporal properties of neuronal populations (Wang, 1999; Chaudhuri et al., 2015; Murray et al., 2018; Wang, 2020). At the circuit level, features such as recurrent connectivity and the balance of excitation and inhibition (E/I balance) strongly shape local dynamics (Chaudhuri et al., 2015; Wang, 1999; 2002; 2008; 2020; Murray et al., 2018). Densely recurrent, feedback-rich circuits can sustain activity over longer periods (yielding longer timescales), whereas predominantly feedforward architectures promote rapid signal decay and shorter timescales. Specifically, strong recurrent excitation, mediated by slow NMDA receptors (Wang, 1999), is thought to maintain the persistent and sustained activity in the absence of immediate sensory stimulation (Wang, 2008; 2020; Chaudhuri et al., 2015; Gao et al., 2020). These properties are not uniformly distributed across the cortex but instead vary systematically along cortical hierarchies, constraining the temporal integration properties of neural populations (Murray et al., 2018; Beul & Hilgetag, 2019). Higher-order association regions generally exhibit longer integration windows than primary sensory areas that have more transient and rapid changes in activity in response to immediate external stimuli (Murray et al., 2014; Shafiei et al., 2020). The sustained activity and longer integration windows observed in association cortex are potentially supported by a combination of local recurrent and feedback circuitry as well as broader system-level interactions that stabilize persistent activity (Wang, 2020; Kauvar et al., 2025).

At the systems level, intrinsic timescales have been found to provide robust and interpretable measures of spontaneous neural dynamics across modalities (e.g. fMRI, ECoG/EEG, single-unit recordings; Watanabe et al., 2019; Gao et al., 2020; Golesorkhi et al., 2021a). Notably, convergent results across modalities support the idea that intrinsic timescale captures a conserved aspect of cortical dynamics. In humans, resting-state fMRI has made intrinsic timescale particularly accessible for mapping these dynamics at the whole-brain level. Previous studies have identified links between measures of intrinsic timescale and cognition and behavior (Watanabe et al., 2019; Gao et al., 2020; Golesorkhi et al., 2021b; Wang, 2022), as well as their implications in brain disorders and diseases (Wengler et al., 2020; Xie et al., 2023). However, less is known about how these dynamics change over development.

Intrinsic timescale can be quantified in several ways, including lag-1 temporal autocorrelation, decay time to a fixed threshold of the autocorrelation function (ACF), exponential decay constants, spectral parameterization, and the sum of the positive portion of the ACF (Gao et al., 2020; Raut et al., 2020; Shinn et al., 2023; Bero et al., 2026). Here, we focus on the ACF-sum measure because it is a widely used approach in fMRI studies of intrinsic timescale and incorporates autocorrelation structure across multiple positive lags, rather than relying on a single lag estimate or fixed threshold. This provides a complementary perspective on temporal persistence by capturing both short-lag autocorrelation and the longer positive tail of the ACF. Despite its use in fMRI studies of neural dynamics, relatively little is known about how ACF-sum intrinsic timescale varies across the cortex during youth. Applying this measure in large developmental fMRI datasets therefore fills an important gap and enables comparison with prior fMRI studies using similar full-ACF approaches.

Understanding how the intrinsic timescale develops is crucial for several reasons. First, it sheds light on fundamental aspects of developmental neurobiology, particularly the relationship between processing speeds and cognitive performance (Waschke et al., 2021). Acquiring the appropriate processing speed at different developmental stages is essential for the brain’s optimal function, as varying timescales facilitate distinct types of information processing. For example, fast timescales enable efficient transmission of simple information, while slower timescales support the integration of complex, feedback-driven signals (Chaudhuri et al., 2015; Wang, 2008; 2020). This differentiation is vital for cognitive functions such as memory and decision-making (Gao et al., 2020). Furthermore, recent reports indicate that neural dynamics shift across the lifespan (Gao et al., 2020; Truzzi & Cusack, 2023; Wu & Gollo, 2025; Bero et al., 2026), reinforcing the need to explore how these dynamics change with age and how they relate to cognition and disorders. Additionally, observed alterations in intrinsic timescale in developmental disorders, such as autism spectrum disorder, underscore the relevance of characterizing these developmental patterns (Watanabe et al., 2019). By elucidating how intrinsic timescale matures across childhood and adolescence, we can gain insights into both typical and atypical cognitive development.

However, relatively few studies have examined how intrinsic timescale develops across the cortex in youth. Prior developmental work has primarily focused on electrophysiological modalities such as EEG and ECoG (McKeon et al., 2025; Miles et al., 2026), reporting an overall decrease in intrinsic timescale with age. In contrast, fMRI-based investigations in youth remain limited. One study reported a narrowing of intrinsic timescale in the hippocampus, focusing on a specific structure rather than the whole cortex (Zeng et al., 2024). Another study in infants showed that neonates exhibit a markedly different timescale organization than adults, characterized by generally longer timescales and network-specific variability (Truzzi & Cusack, 2023). Notably, this work also demonstrated that the relative ordering of unimodal and transmodal networks differs between infancy and adulthood, suggesting that development involves a redistribution of timescales across cortex rather than a uniform global shift. Despite these observations, the development of fMRI-based intrinsic timescale across the whole cortex in youth remains relatively uncharacterized.

In this study, we map developmental patterns of intrinsic timescale across childhood and adolescence using resting-state fMRI from a large cohort of typically developing youth. By applying an ACF-sum measure of intrinsic timescale, we characterize temporal persistence across the positive portion of the ACF and evaluate how these regional patterns relate to the cortical hierarchy defined by the S–A axis. We further assess the generalizability of the observed developmental patterns in a large independent developmental sample. We then assess the specificity of our developmental results by also examining associations between intrinsic timescale and age in young adulthood. Together, these analyses clarify how fMRI-based intrinsic timescale varies across youth and how these age-related differences relate to large-scale cortical organization.

## Results

Cortical maps of intrinsic timescale were estimated using the autocorrelation function of resting-state fMRI time series for two cross-sectional, developmental cohorts (ages 8–22 years) and a young-adult dataset (ages 22–37 years) (**Figure 1**). We then used generalized additive models (GAMs) to examine linear and nonlinear associations between fMRI intrinsic timescale and age. We also assessed whether age-related differences in intrinsic timescales align with the cortical hierarchy defined by the S–A axis.

**Figure 1.**
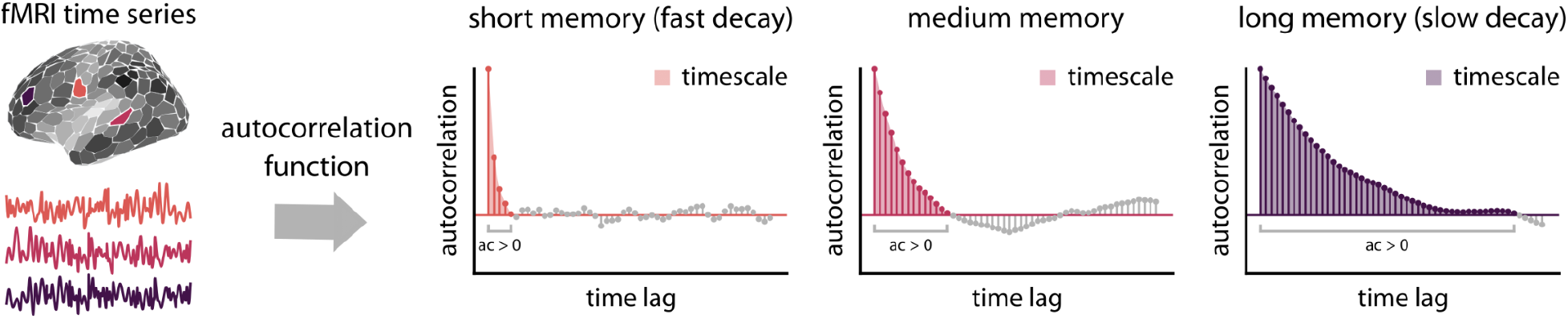
Quantifying intrinsic timescale using autocorrelation function. Fully processed parcellated fMRI time-series were used to estimate intrinsic timescale for each brain region (i.e., parcel) and participant. We estimated an autocorrelation function (ACF) for the normalized (i.e., *z*-scored) time series. Intrinsic timescale was then quantified as the sum of positive autocorrelation values (ac > 0; colored parts of the ACFs above) multiplied by TR for each cortical region and individual. This is approximately equivalent to calculating the area under the curve of the positive part of the ACF curve (i.e., shaded areas). As demonstrated in the example plots, a fast-decaying ACF will have shorter memory, and thus a shorter timescale (orange region and ACF plot), compared to a slow decaying ACF (purple region and ACF plot). Data and code needed to regenerate this figure can be found at https://doi.org/10.5281/zenodo.22799416.

### Intrinsic timescale is longer in association cortex compared to sensorimotor cortex in youth

We first sought to characterize intrinsic timescale patterns in the two developmental cohorts. Intrinsic timescale varied across the cortex in both developmental datasets: the Human Connectome Project–Development (HCPD; *n*=565; Somerville et al., 2018) and the Healthy Brain Network (HBN; *n*=729; Alexander et al., 2017) (**Figure 2**). Visual inspection suggested that the cortical distribution of fMRI intrinsic timescale was broadly segregated into lower-order unimodal regions and higher-order transmodal areas (**Figure 2a**). The mean timescale maps, averaged across individuals, were positively correlated between the two developmental datasets, demonstrating consistent patterns (**Figure 2a**; *r*_s_ = 0.76, *p*_spin_ = 0.0002). To assess whether timescale maps in youth reflect the cortical hierarchy, we directly compared the average timescale to the S–A axis (Sydnor et al., 2021; **Figure 2b**). The cortical distribution of timescale was positively associated with the S–A axis in both datasets. However, this association was statistically significant only in HCPD (**Figure 2b** scatter plots: HCPD: *r*_s_ = 0.37, *p*_spin_ = 0.03; HBN: *r*_s_ = 0.27, *p*_spin_ = 0.12). Further analysis demonstrated that intrinsic timescale was generally significantly longer in association cortex compared to sensorimotor cortex in both datasets (**Figure 2b** boxplots: HCPD: *t* = 7.8, *p*_spin_ = 0.01; HBN: *t* = 6.6, *p*_spin_ = 0.04; two-tailed). Collectively, these findings indicate that the intrinsic timescale tends to be longer in the association cortex compared to the sensorimotor cortex in youth.

**Figure 2.**
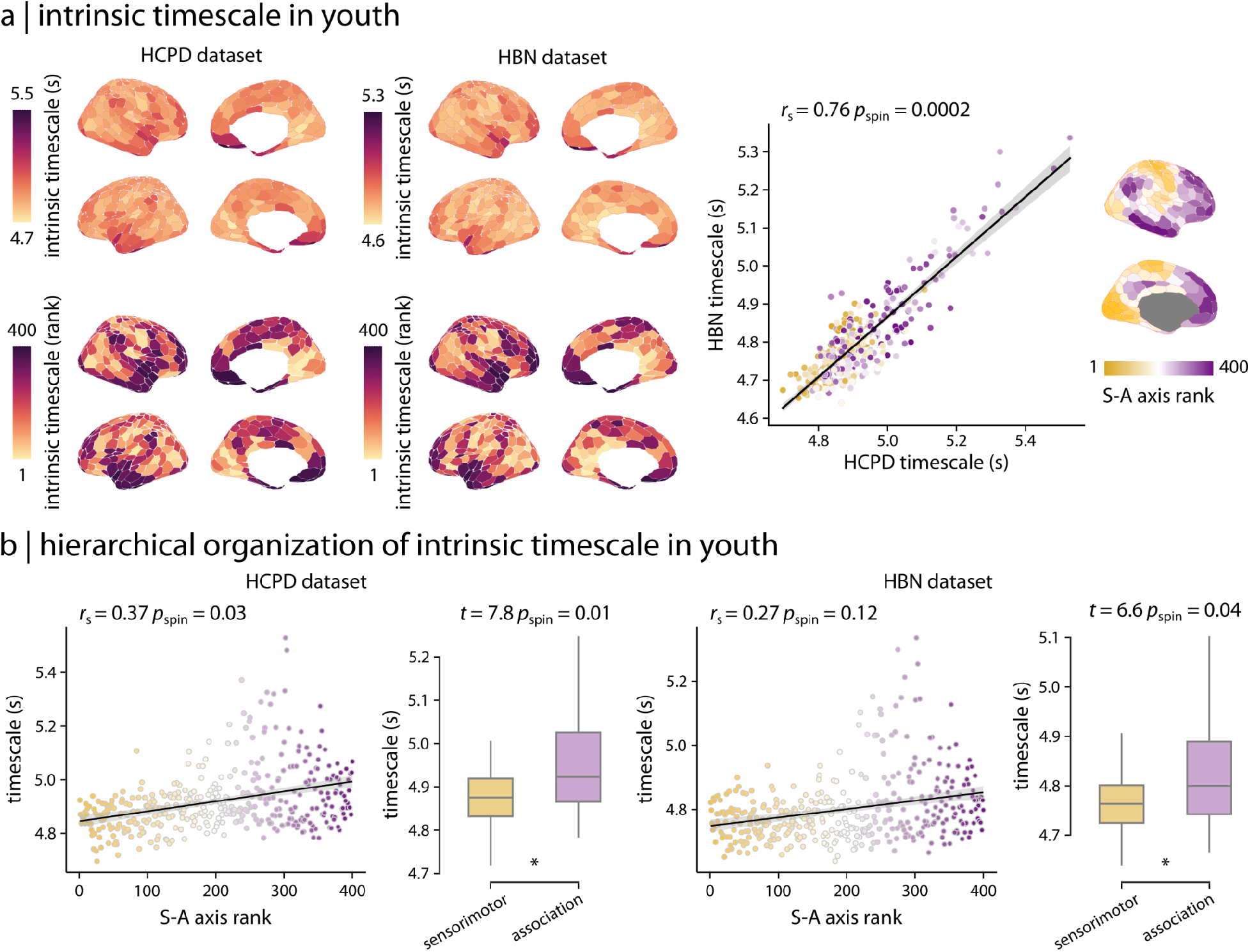
Intrinsic timescale is longer in association cortex compared to sensorimotor cortex in youth. **(a)** Cortical distribution of intrinsic timescale reflects similar patterns in two independent developmental cohorts: the HCPD and HBN datasets. Intrinsic timescale maps were compared directly between the two datasets (scatter plot). Each brain region (i.e., each circle in the scatter plot) is colored based on the region’s rank along an organizational axis of the cortex that spans sensorimotor to association cortices (S–A axis). **(b)** Intrinsic timescale in youth (HCPD and HBN dataset) reflects the hierarchical organization captured by the S–A axis, such that sensorimotor regions have shorter timescales while association regions have longer timescales. Significance of the associations between cortical maps (i.e., scatter plots) was assessed using spatial autocorrelation-preserving permutation tests (i.e., spin test). *r*_s_ denotes Spearman’s rank correlation coefficient. Linear regression lines are added for visualization purposes only. Asterisks in the boxplot denote significant differences in the means (two-tailed *t*-test). Data and code needed to regenerate this figure can be found at https://doi.org/10.5281/zenodo.22799416.

### Neurodevelopmental patterns in intrinsic timescale align with the cortical hierarchy

To characterize the developmental patterns of intrinsic timescale in youth, we examined the relationship between participant age and both whole-brain and regional timescale in HCPD, while controlling for participant sex and in-scanner motion. We found that the whole-brain average timescale increases during development in youth (**Figure 3a**; partial *R*^2^ = 0.047, *p*_anova_ = 1.25 x 10^-7^). Furthermore, we found that regional neurodevelopmental patterns were heterogeneous across the cortex, with smaller age effects in the sensorimotor cortex and larger age effects in association cortex (**Figure 3b**). Notably, the effect size of the developmental associations aligned with the cortical hierarchy defined by the S–A axis (**Figure 3c**; *r*_s_ = 0.41, *p*_spin_ = 0.001). Consistent with this hierarchical pattern, supplementary within-subject analyses showed that the categorical separation between sensorimotor and association timescales increased with age (**Figure S1**). Further inspection of regional developmental model fits revealed that timescale increased in association cortex but was stable in sensorimotor regions (**Figure 3d**). Overall, these findings demonstrate that fMRI intrinsic timescale increases with age in youth, mainly driven by increases in timescale in association cortex. Timescale development reflected a hierarchical pattern, recapitulating the S–A axis of cortical organization.

**Figure 3.**
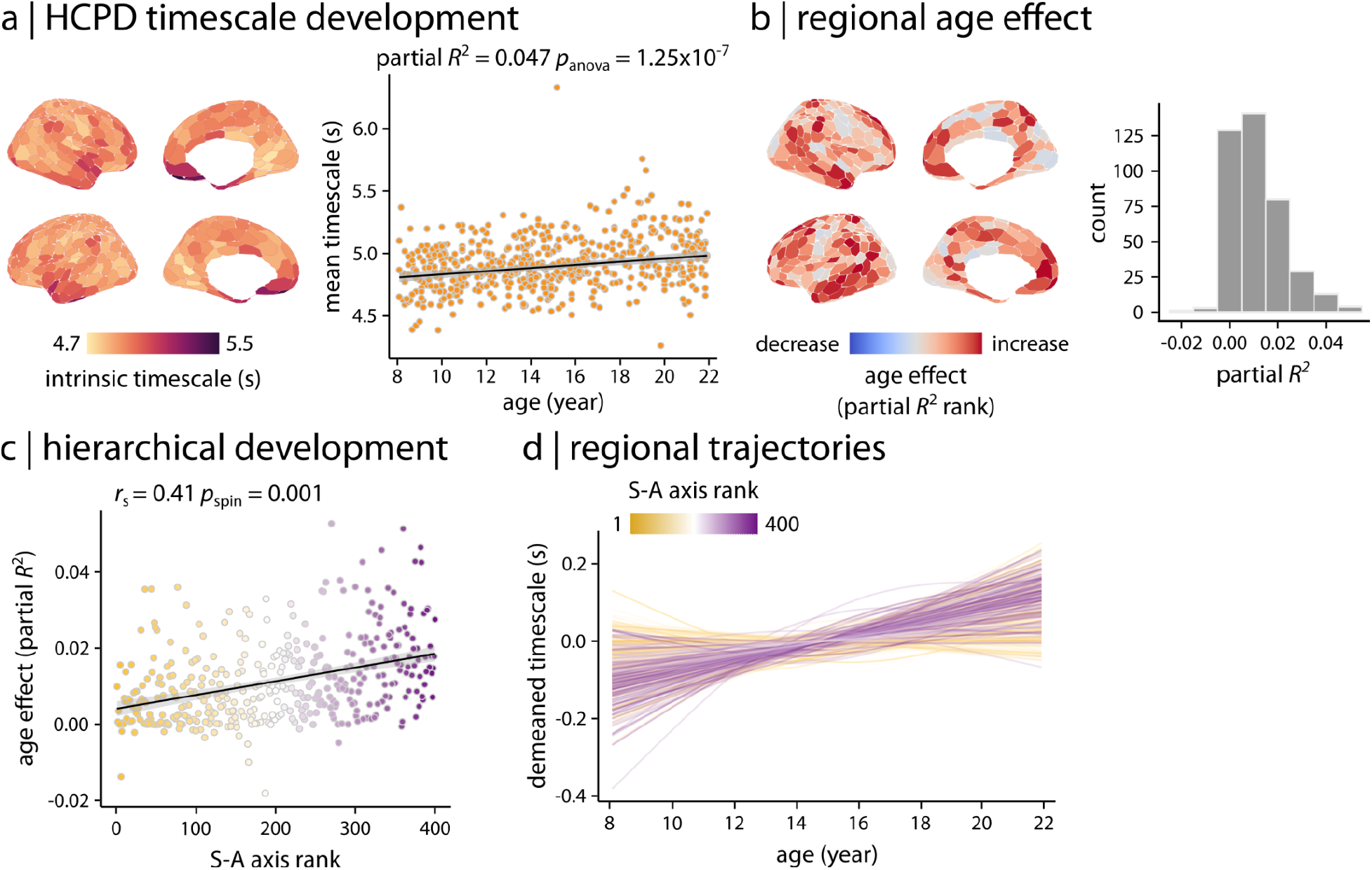
Developmental patterns of intrinsic timescale. Generalized additive models (GAMs) were used to assess linear and nonlinear age effects in intrinsic timescale. **(a)** A whole-brain GAM was applied to model age-related differences in cortex-wide average timescale. GAM results demonstrated that average timescale increases during development in youth (partial *R*^2^ = 0.047, *p*_anova_ = 1.25 x 10^-7^). **(b)** Region-wise GAMs were used to examine developmental patterns in intrinsic timescale at the regional level. Age effects quantified as partial *R*^2^ are depicted across the cortex, displaying a heterogeneous spatial distribution. **(c)** Age effects on intrinsic timescale were compared to the S–A axis, identifying a hierarchical pattern of developmental patterns in intrinsic timescale along the S–A axis. Significance of the association between age effects and S–A axis rank was assessed using 10,000 spin tests. *r*_s_ denotes Spearman’s rank correlation coefficient. Linear regression line is added for visualization purposes only. **(d)** Regional trajectories of developmental patterns in intrinsic timescale were obtained from region-wise GAM results (as shown in panel **b**). Each line corresponds to the model fit of each cortical region and is colored based on the region’s rank along the S–A axis. The results demonstrate that intrinsic timescale in association regions increases during development in youth while it remains relatively stable in sensorimotor regions. Data and code needed to regenerate this figure can be found at https://doi.org/10.5281/zenodo.22799416.

### Findings replicate in an independent developmental cohort

We next examined the extent to which these developmental findings were generalizable by replicating our analyses in the HBN dataset. Consistent with HCPD, we found that the average intrinsic timescale increased during development in HBN (**Figure 4a**; partial *R*^2^ = 0.037, *p*_anova_ = 1.13 x 10^-7^). Regional analysis similarly identified heterogeneous developmental effects across the cortex (**Figure 4b**). As in HCPD, these developmental effects were aligned with the S–A axis (**Figure 4c**; *r*_s_ = 0.19, *p*_spin_ = 0.02). Developmental fits showed that overall, timescale remained stable in sensorimotor regions but increased in association cortex (**Figure 4d**). To directly evaluate the degree to which results converged across the two datasets, we compared developmental patterns of intrinsic timescale between HBN and HCPD (**Figure 5**). We found that maps of associations with age (i.e., partial *R*^2^ maps) were similar for the two datasets (**Figure 5**; *r*_s_ = 0.30, *p*_spin_ = 0.0002). Overall, 46.5% of the regions in HCPD and 22.5% of the regions in HBN demonstrated significant age effects after FDR correction. The developmental patterns in these regions had a significant spatial overlap between HCPD and HBN (**Figure 5**; Dice score = 0.42, *p*_spin_ = 0.0008). For regions with at least one significant derivative, we summarized the timing of maximal development as the age at which the regional model derivative reached its maximum absolute value (**Figure S2a**). These estimates should be interpreted cautiously because age of maximal development is only defined for regions with significant derivative-based change; regions with weak, stable, or uncertain age trajectories do not have a well-defined peak-change age. To complement this analysis, we also examined the age at which whole-cortex rates of timescale differences were maximally aligned with the S-A axis, which peaked in early-to-mid adolescence in both datasets (**Figure S2b**; 14.6 years in HCPD and 12.8 years in HBN).

**Figure 4.**
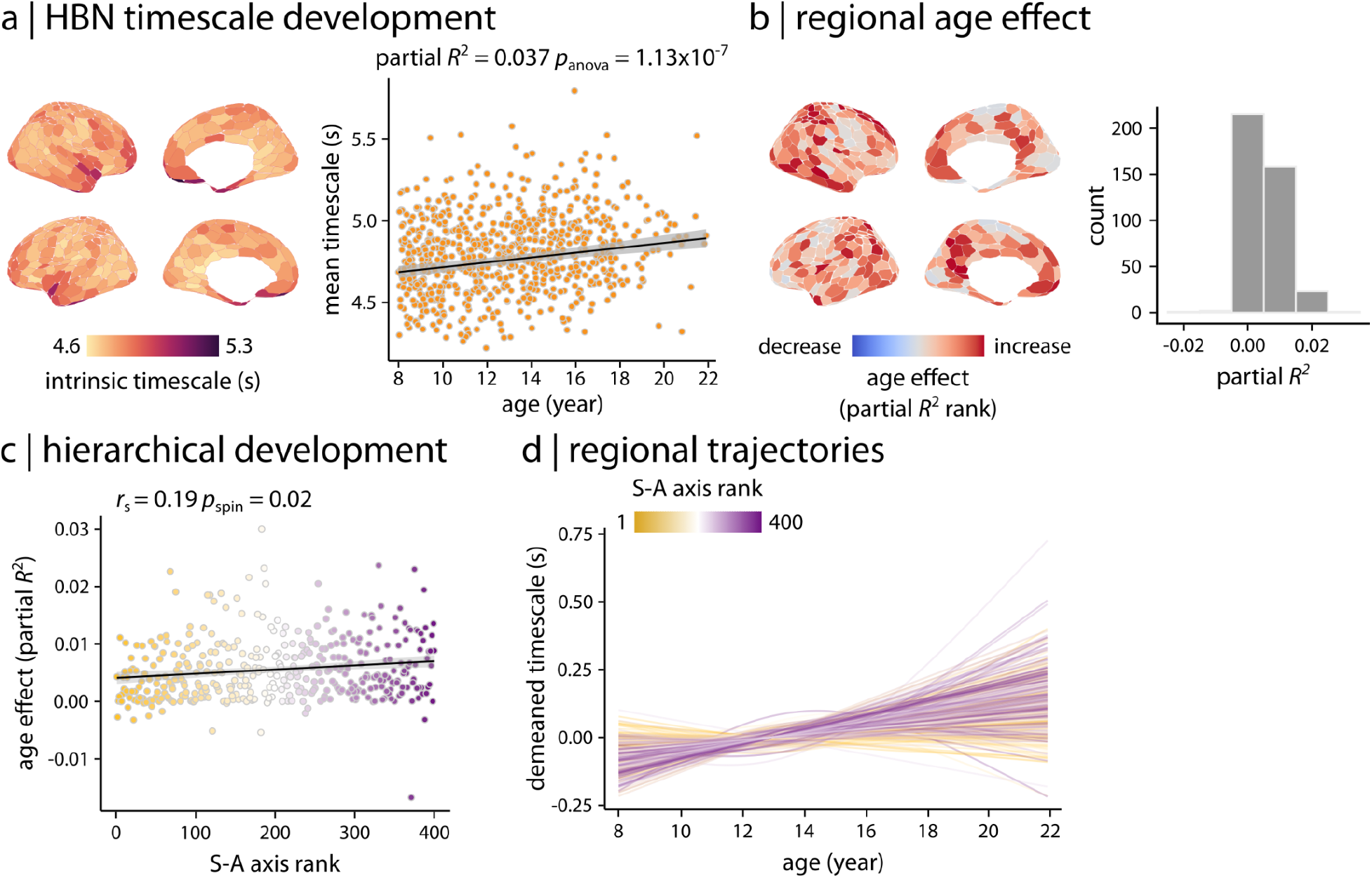
Replication in an independent dataset. Developmental patterns of intrinsic timescale were replicated in an independent dataset (i.e., HBN). **(a)** Consistent with the finding in the HCPD dataset, GAM results demonstrated that the average timescale increases during development in youth in the HBN dataset (partial *R*^2^ = 0.037, *p*_anova_ = 1.13 x 10^-7^). **(b)** Region-wise GAMs identified heterogeneous age effects (i.e., partial *R*^2^) across the cortex. **(c)** Age effects on intrinsic timescale were hierarchically organized along the S–A axis. Significance of the association between age effects and S–A axis rank was assessed using 10,000 spin tests. *r*_s_ denotes Spearman’s rank correlation coefficient. Linear regression line is added for visualization purposes only. **(d)** Regional trajectories of developmental patterns in intrinsic timescale were obtained from region-wise GAM results (as shown in panel **b**). Each line corresponds to the model fit of each cortical region and is colored based on the region’s rank along the S–A axis. Similar to HCPD, the results demonstrate that intrinsic timescale in association regions increases during development in youth while it remains relatively stable in sensorimotor regions. Data and code needed to regenerate this figure can be found at https://doi.org/10.5281/zenodo.22799416.

**Figure 5.**
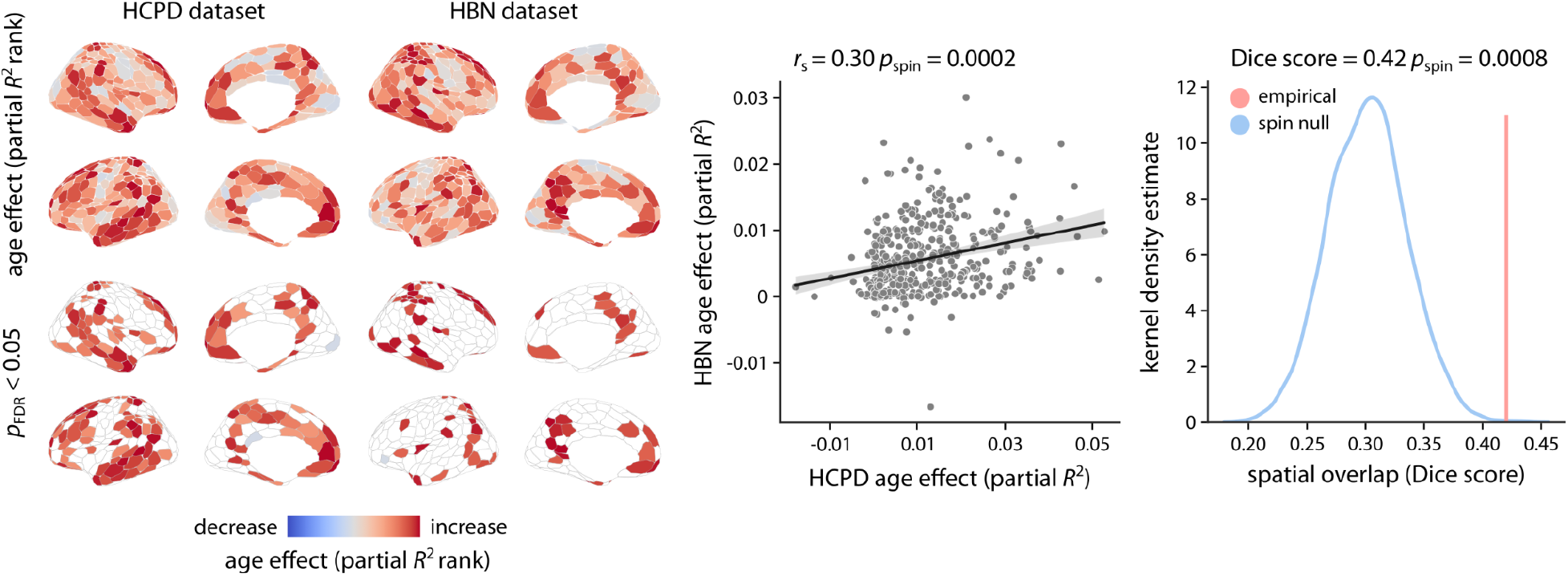
Consistent age effects across two developmental cohorts. Age effects (i.e., partial *R*^2^) obtained from the HCPD and HBN dataset were directly compared (scatter plot). Significance of the association between age effects from the two datasets was assessed using 10,000 spatial autocorrelation-preserving permutation tests (i.e., spin test). *r*_s_ denotes Spearman’s rank correlation coefficient. Linear regression line is added for visualization purposes only. Additionally, the spatial overlap between regions with significant age effects after FDR correction was quantified using Dice score. To assess the statistical significance of the between-dataset spatial overlap, spin tests were used to generate a null distribution of Dice scores (10,000 repetitions). The empirical Dice score was then compared to the null distribution of scores to calculate a *p*-value for the spatial overlap. Data and code needed to regenerate this figure can be found at https://doi.org/10.5281/zenodo.22799416.

### Timescale remains relatively stable in adulthood

To assess whether the observed age-related differences in timescale were specific to childhood and adolescence or extended into young adulthood, we repeated all analyses in an independent sample of young adults (i.e., 22–37 years old) from the Human Connectome Project–Young Adults (HCPYA; van Essen et al., 2013; *n*=973) (**Figure 6**). As expected, intrinsic timescale in young adults reflected the hierarchical organization captured by the S–A axis (**Figure 6a**; *r*_s_ = 0.4, *p*_spin_ = 0.01), such that sensorimotor regions had shorter timescales while association regions had longer timescales (*t* = 7.5, *p*_spin_ = 0.02, two-tailed). However, unlike the developmental findings, age analysis demonstrated that the whole-brain average timescale remains relatively stable in adulthood (**Figure 6b**; partial *R*^2^ = −0.004, *p*_anova_ = 0.04). Regional age effects estimated using GAMs were not statistically significant for the majority of brain regions (11.7% of regions with significant age effects; **Figure 6c**) and did not exhibit significant variability along the S–A axis (**Figure 6d**; *r*_s_ = −0.1, *p*_spin_ = 0.58). Overall, these results emphasize the specificity of the developmental findings in HCPD and HBN to the developmental period of 8 to 22 years old, suggesting that fMRI intrinsic timescale develops along the S–A axis in youth and stabilizes in adulthood. Given the differences in acquisition parameters and cohort characteristics, the young adult dataset was used to assess the specificity of the developmental findings to the developmental cohorts and the stability of age-related effects within young adulthood, rather than as a direct continuation of the developmental age trajectories.

**Figure 6.**
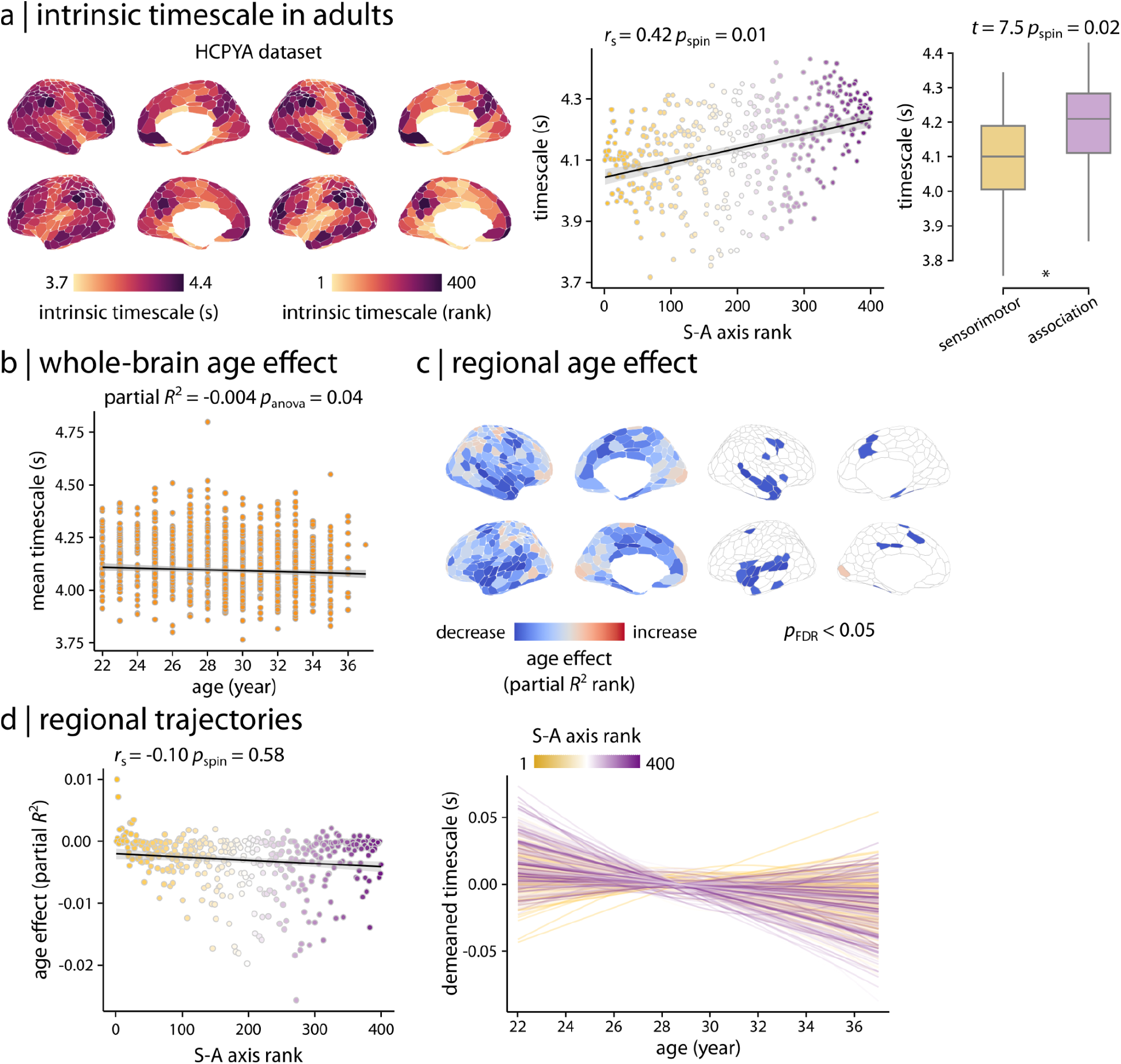
Stable intrinsic timescale in young adults. **(a)** Intrinsic timescale in young adults (HCPYA dataset) reflects the hierarchical organization captured by the S–A axis, such that sensorimotor regions have shorter timescales while association regions have longer timescales. Asterisk in the boxplot denotes significant difference in the means (two-tailed *t*-test). **(b)** However, unlike the developmental cohorts, GAM results demonstrated that the whole-brain average timescale remains relatively stable in adulthood (partial *R*^2^ = −0.004, *p*_anova_ = 0.04). **(c)** Region-wise GAMs identified heterogeneous age effects (i.e., partial *R*^2^) across the cortex; however, those age effects were not statistically significant for the majority of brain regions. **(d)** Unlike the developmental cohorts, age effects on the intrinsic timescale did not reflect the hierarchical organization along the S–A axis during adulthood. Significance of the associations between cortical maps (i.e., scatter plots) was assessed using 10,000 spin tests. *r*_s_ denotes Spearman’s rank correlation coefficient. Linear regression lines are added for visualization purposes only. Data and code needed to regenerate this figure can be found at https://doi.org/10.5281/zenodo.22799416.

### Sensitivity analysis

To evaluate whether our findings were influenced by confounding factors and specific analytical choices, we performed a series of sensitivity analyses. We first examined whether the findings were independent from confounding factors such as brain volume and parcel size. Given that brain size changes during development, total brain volume might impact the average whole-brain intrinsic timescale. However, we only found a weak association between estimated Total Intracranial Volume (eTIV) and intrinsic timescale (**Figure S3a**). To ensure that the developmental patterns of intrinsic timescale were independent from this weak correlation, we repeated the age analysis while including eTIV as a model covariate. The results were consistent with the original findings (**Figure S3b**). Additionally, given that parcel-wise time series are calculated as the average time series across vertices within a given parcel, parcel size might directly impact smoothness of parcel time series and their autocorrelation functions. To test this, we compared parcel size and intrinsic timescale for each parcel and each participant in HCPD and found no associations (mean *r*_s_ = 0.01). Moreover, given that HBN is a more heterogeneous sample compared to HCPD and includes help-seeking individuals with higher levels of psychopathology, we repeated all analyses using two subsets of HBN individuals (each with *N*=600; this approximately matches the HCPD sample size) with low motion or low psychopathology (i.e., *p*-factor). The findings were consistent with the full HBN sample (**Figure S4**). Furthermore, to assess the extent to which the results were influenced by the choice of atlas and number of parcels, we repeated the analysis with a lower resolution Schaefer atlas with 200 parcels in HCPD. The results were consistent with the original findings with the Schaefer 400 atlas (**Figure S5**). Moreover, to ensure that findings were independent from signal-to-noise ratio (SNR) of the fMRI signal, we estimated temporal SNR (tSNR) as the ratio of the time-series mean to standard deviation for each region and participant in HCPD. Neither the intrinsic timescale nor the age effects (partial *R*^2^) were significantly associated with tSNR (**Figure S6**). Additionally, to ensure that findings were independent from data acquisition sites for HCPD and HBN, we repeated the analyses after harmonizing intrinsic timescale across sites. Findings were almost perfectly correlated before and after harmonization for both HCPD and HBN datasets (**Figure S7**).

To test whether intrinsic timescale estimates were sensitive to methodological choices and to better contextualize our findings with prior work, we compared our ACF-sum measure with several alternative approaches for quantifying intrinsic timescale in HCPD. First, we quantified intrinsic timescale using two other approaches based on full ACF: (1) fitting an exponential function to ACF and using the exponential decay constant to estimate intrinsic timescale (Ito et al., 2020); (2) directly using the first zero-crossing point of ACF as intrinsic timescale. Both approaches generated intrinsic timescale maps and developmental patterns consistent with the original method (**Figure S8**). We further repeated the HCPD analyses using lag-1 temporal autocorrelation (lag-1 TA), following prior work (Shinn et al., 2023; Bero et al., 2026). We found that lag-1 TA and ACF-sum timescale differed in their mean spatial distribution but showed partially overlapping developmental effects (**Figures S9-S10**). We also decomposed the ACF-sum timescale into short- and long-lag components and found that lag-1 TA was most closely related to short-lag autocorrelation, whereas the full ACF-sum measure was more strongly influenced by longer-lag positive autocorrelation (**Figure S11**). Together, these analyses suggest that our core findings are robust across several full-ACF-based timescale definitions, while also showing that different metrics emphasize different features of temporal persistence: lag-1 TA primarily reflects short-lag autocorrelation, whereas the ACF-sum measure is more sensitive to longer-lag positive autocorrelation across the full ACF. This suggests that the developmental effects observed with our ACF-sum measure were driven primarily by long-lag ACF structure.

## Discussion

We systematically characterized how intrinsic timescale measured using fMRI develops during youth. We found that intrinsic timescales are generally longer in the association cortex and shorter in the sensorimotor cortex in youth. Furthermore, modeling the regional developmental trajectories of timescale demonstrated that intrinsic timescale development aligned with the cortical hierarchy. Importantly, these developmental patterns were robust, appearing in two large independent developmental cohorts. Moreover, findings in a young adult sample demonstrated that age-related differences in the intrinsic timescale are specific to the developmental cohorts and that the intrinsic timescale appears to remain stable in young adulthood. Together, these findings position intrinsic timescale as an fMRI measure sensitive to neurodevelopment in youth that helps bridge levels of analysis, linking local circuit dynamics to the brain’s broader hierarchical maturational program.

The observed S–A axis pattern of timescale development aligns with well-established principles of cortical organization. In adults, neuronal timescales are known to increase in length from primary sensorimotor regions to higher-order association cortices, mirroring fundamental anatomical and functional gradients (Murray et al., 2014; Demirtas et al., 2019; Gao et al., 2020; Ito et al., 2020). Recent developmental studies have suggested that similar large-scale cortical axes guide maturation (Sydnor et al., 2021; Luo et al., 2024; Nishio et al., 2024; Bero et al., 2026). Our findings suggest that age-related differences in fMRI intrinsic timescale are broadly organized along the cortical hierarchy. Across youth, regional patterns of intrinsic timescale differences are aligned with the sensorimotor-to-association gradient that defines adult brain architecture. Additionally, supplementary within-subject analyses further suggest that the categorical separation between sensorimotor and association timescales also increases with age. Our results also mirror other metrics of brain functional development. Notably, previous work has shown that the maturation of both spontaneous activity amplitude (i.e., ALFF) (Sydnor et al., 2023) and functional connectivity (Luo et al., 2024) unfold along the S–A axis, indicating a hierarchical refinement of neural activity during youth.

While this work was being prepared, a very recent study reported that age-related differences in temporal scales of resting-state cortical activity reflect maturation of cortical myelination during the human lifespan and align with the S–A axis (Bero et al., 2026). In this study, the temporal scales were quantified using lag-1 temporal autocorrelation (lag-1 TA) of fMRI time series using data from HCPD. Bero et al. (2026) also demonstrated that age-related differences in temporal scales vary across the cortex, such that the temporal scales decrease the most in sensorimotor regions (e.g., visual and somatomotor cortices) while they increase in prefrontal areas (Bero et al., 2026). Our findings provide convergent evidence for how neural dynamics develop in youth. Although our measure of intrinsic timescale conceptually overlaps with the temporal scales defined by Bero et al. (2026), the two measures characterize neural dynamics in different ways. Specifically, the measure of timescale we used takes into account the full shape of the autocorrelation function of fMRI time series, whereas the measure of temporal scales in Bero et al. (lag-1 TA) mainly focuses on the autocorrelation function at short lags (i.e., lag 1). Despite these differences in how fMRI dynamics were defined, both studies converge in showing that the development of neural dynamics is spatially heterogeneous across cortex and broadly related to large-scale cortical hierarchy. Additionally, the regional heterogeneity of age-related differences observed across both measures suggests that they capture partially overlapping but distinct aspects of neural dynamics. For example, we found that intrinsic timescales remain stable in sensorimotor regions while they increase in association cortex, whereas Bero et al. found that temporal scales decrease in sensorimotor regions while they increase in prefrontal regions.

To further investigate these differences, we repeated our developmental analyses using lag-1 TA in the same HCPD sample and preprocessing pipeline used for our primary ACF-sum analyses. This comparison showed that lag-1 TA and ACF-sum timescale had distinct regional mean maps but demonstrated partially overlapping regional age effects. Decomposing the ACF-sum timescale into short-and long-lag components further suggested that lag-1 TA and ACF-sum timescale emphasize partially distinct features of the ACF: lag-1 TA primarily reflects immediate short-lag decorrelation, whereas ACF-sum timescale is more sensitive to the broader shape of the ACF, including sustained positive autocorrelation across longer lags. Thus, while methodological differences in preprocessing, sample selection, and modeling may contribute to discrepancies across studies, our results indicate that some differences may also arise because commonly used timescale metrics emphasize different characteristics of the same fMRI signal.

These subtle but important differences underscore the need to integrate multiple approaches to quantifying neural dynamics across imaging modalities and datasets to obtain a more comprehensive view of brain development in youth. For example, prior research in infants suggests that neonates exhibit a distinct timescale organization compared to adults, characterized by generally longer timescales and network-specific variability (Truzzi & Cusack, 2023). Although this study used the same measure of intrinsic timescale as ours, the age range differed substantially (infancy vs. childhood and adolescence). While longer timescales in neonates may relate to slower baseline information processing, they may also reflect differences in how information is integrated in early development. Notably, this work shows that the relative ordering between unimodal and transmodal cortices differs between infancy and adulthood (i.e., higher timescales in unimodal regions in infancy vs. transmodal regions in adulthood). Reconciling these findings with our results therefore likely requires developmental shifts in the relative ordering of timescales across cortical systems.

Furthermore, prior studies using imaging modalities with higher temporal resolution, such as EEG and ECoG, have reported overall decreases in intrinsic timescale during development (McKeon et al., 2025; Miles et al., 2026). Although similar measures of timescale were used across these studies and our study in fMRI, the overall age-related trends in timescale varied across modalities. These differences may reflect the fact that different imaging modalities probe distinct aspects of neural dynamics, as they are sensitive to different frequency ranges and may preferentially capture activity arising from different laminar sources (Scheeringa et al., 2016; Sadaghiani & Wirsich, 2020). In this context, developmental decreases in electrophysiological timescale and increases in fMRI-based ACF-sum timescale may not necessarily reflect opposing developmental patterns. Instead, electrophysiological studies may be more sensitive to faster local neural dynamics, whereas the present fMRI-based measure reflects the persistence of slow BOLD fluctuations within the 0.01-0.08 Hz range. Because temporal filtering influences the shape of the autocorrelation function (Shumway & Stoffer, 2017), restricting the signal to slower BOLD fluctuations can produce a more slowly decaying ACF. Consistent with this interpretation, our short- and long-lag decomposition of the fMRI autocorrelation function suggests that the ACF-sum measure is particularly sensitive to longer-lag positive autocorrelation, indicating that the association-cortex increases observed here should not be interpreted as a global slowing of neural activity across all temporal scales. Finally, discrepancies across studies, even within the same modality, may also arise from differences in data acquisition, preprocessing pipelines, and analytic approaches. Such methodological variation can influence estimates of resting-state dynamics (e.g., time series properties, cross-correlations, and intrinsic timescale; Li et al., 2024) and should be considered when interpreting findings.

We found that the intrinsic timescale increases with age in youth, particularly in the association cortex. This suggests that neural dynamics in association cortices become more integrative as the brain matures. While faster dynamics may facilitate immediate responses, longer timescales in association cortex are likely critical for supporting higher-order cognitive functions such as working memory (Gao et al., 2020). Convergence with established cortical gradients further indicates that intrinsic timescale follows a common developmental program aligned with the brain’s hierarchical organization. By charting normative timescale development from mid-childhood through adolescence, our study helps contextualize findings from both earlier and atypical development (Truzzi & Cusack, 2023; Watanabe et al., 2019). Specifically, our results suggest that from middle childhood onward, intrinsic timescales adopt an adult-like hierarchical ordering, helping bridge the gap between neonatal and mature states. This normative trajectory is also relevant for understanding neurodevelopmental disorders. For example, autism spectrum disorder has been associated with atypical timescale development, where the typical variation in timescales is disrupted (Watanabe et al., 2019). Establishing how timescales mature in typically developing youth therefore provides an important baseline against which such deviations can be interpreted.

The present findings must be interpreted in the context of several methodological considerations. First, our analysis is cross-sectional, which limits inferences about within-person developmental trajectories. Longitudinal studies will be needed to confirm how intrinsic timescales evolve in the same individuals over time and to assess whether individual differences in timescale trajectories predict neurocognitive outcomes or vulnerability to mental health conditions. Second, our analysis includes childhood, adolescence, and early adulthood (age range of 8–22 years), missing critical information about the developmental patterns of intrinsic timescale in infancy and early childhood. Future studies should seek to combine data collected with similar acquisition parameters across a broader range of ages in development to evaluate this possibility directly. Third, the intrinsic timescale was measured via fMRI signals with a moderate sampling rate (TR=0.8s), which constrains the temporal resolution. Faster neural events or fine-grained timescale differences might be missed due to the coarse temporal sampling of fMRI. Future multimodal studies are required to address the cross-modal correspondence of developmental patterns in neural dynamics. Finally, we note that our focus was on cortical timescales at the regional level. However, neural dynamics and their characteristics might vary systematically between different brain structures (Zeng et al., 2024). Developmental patterns in fMRI intrinsic timescale remain to be examined in subcortical structures and using finer spatial gradients.

Taken together, our findings suggest that development of intrinsic timescale aligns with the cortical hierarchy, with an increase in timescale in the association cortex that likely reflects an expansion of integrative capacity. These results open new avenues for investigating the mechanistic underpinnings of timescale development. Moving forward, it will be valuable to link these findings to multimodal data and evaluate timescale in the context of developmental processes like synaptic pruning, myelination, or shifts in excitation-inhibition balance. Such work could help determine why association cortices exhibit prolonged integration windows during youth while sensorimotor areas remain stable, providing biological insight into the hierarchical development we observed.

## Methods

### Participants

We used resting-state functional magnetic resonance imaging (fMRI) data from 3 independent cohorts: (1) a cross-sectional developmental dataset used as a discovery sample; (2) an independent cross-sectional developmental dataset used to replicate the findings; (3) a young adult dataset to evaluate the specificity of our results.

#### Discovery sample – HCPD

The discovery developmental sample included resting-state fMRI data from the Human Connectome Project–Development (HCPD; Somerville et al., 2018; *n*=565, age range 8–22 years, 302 female and 263 male). HCPD is a sample of typically developing children and adolescents recruited in the United States of America across four sites: University of Minnesota, Harvard University, Washington University in St. Louis, and University of California-Los Angeles. All study procedures were approved by a central Institutional Review Board at Washington University in St. Louis.

Imaging data were obtained on 3 Tesla Siemens Prisma platforms at the four sites. All HCPD participants had T1 scans passing quality control (QC). The fMRI data included 26 minutes of resting-state scans per participant in total acquired in four runs with the following acquisition parameters: multiband acquisition, 2mm^3^, TR=800ms, TE=37ms, flip angle=52°, number of volumes=478. We only included fMRI scans with low in-scanner motion (i.e., mean framewise displacement (FD) < 0.2mm; see “Neuromaging data processing” for details); these QC criteria excluded *n*=48 individuals from the initial sample of *n*=635 individuals with resting-state data. Additionally, from the remaining sample of *n*=587 individuals, we excluded *n*=20 individuals with medical conditions resulting in gross neurological abnormalities or affecting brain function (e.g., individuals with cancer or leukemia, lead poisoning, sickle cell anemia, accidental poisoning, multiple sclerosis, seizure, and brain injury). Finally, we limited our sample to the age range of 8–22 years old, resulting in a final sample with *n*=565 individuals.

#### Replication sample – HBN

The replication developmental sample included resting-state fMRI data from the Healthy Brain Network (HBN; Alexander et al., 2017; *n*=729, age range 8–22 years, 279 female, 418 male, 32 other). HBN is a sample of children and adolescents residing in the New York City area (United States of America) that is aimed to represent a diverse sample of healthy and help-seeking individuals. The HBN study was approved by the Chesapeake Institutional Review Board.

Our HBN sample included data acquired at four different acquisition sites: Staten Island (SI - Mobile Scanner), Rutgers University (RU), The City University of New York (CUNY), and Citigroup Biomedical Imaging Center (CBIC). Note that we excluded HBN data from the Staten Island (SI) site from our analysis given that SI’s fMRI data was acquired on a different scanner (1.5T Mobile Scanner) with different acquisition parameters. Image acquisition parameters for functional images collected at the RU, CUNY, and CBIC sites were as follows: multiband acquisition, 2.4mm^3^, TR=800ms, TE=30ms, flip angle=31°, number of volumes=375 (median; over 97% of scans had 375 volumes, with a small subset containing a different number of volumes). We excluded *n*=93 participants who did not pass T1 QC from the initial sample of *n*=1743 with resting-state data (for details, see Shafiei et al., 2025). As for HCPD, we also excluded *n*=831 participants from the remaining sample of *n*=1650 due to in-scanner motion (i.e., mean FD < 0.2mm). Finally, same as in HCPD, we limited our sample to the age range of 8–22 years old, resulting in a final sample with *n*=729 individuals.

#### Young adult sample to evaluate specificity – HCPYA

Resting-state fMRI data from healthy young adults were obtained from the Human Connectome Project–Young Adults (HCPYA; van Essen et al., 2013; *n*=973, age range 22–37 years, 523 female, 450 male). All HCPYA imaging data were collected on a customized 3 Tesla scanner at Washington University (WashU). The study procedures were conducted under Institutional Review Board–approved protocols of the WU–Minn Human Connectome Project consortium (Van Essen et al., 2013).

As for HCPD, all HCPYA participants had T1 scans passing quality control (QC). The fMRI Data included 1 hour of resting-state scans acquired over 2 days (2 scans along opposing R/L and L/R phase encoding directions per day; each scan was approximately 15 minutes long). All functional images were acquired with the following acquisition parameters: multiband acquisition, 2mm^3^, TR=720ms, TE=33.1ms, flip angle=52°, number of volumes=1200 (median; over 99% of scans had 1200 volumes). As for the developmental data, we excluded n=123 participants from the initial sample of n=1097 individuals (with resting-state data) due to in-scanner motion (i.e., mean FD < 0.2mm), resulting in a final sample with *n*=973 individuals.

### Neuroimaging data processing

All three datasets were preprocessed and post-processed with similar image processing pipelines. The detailed descriptions are provided based on reports automatically generated by the implemented processing tools, and are released under the CC0 1.0 Universal license.

#### Developmental sample preprocessing: (HCPD and HBN)

The two developmental datasets (i.e., HCPD and HBN) were preprocessed with fMRIPrep version 22.0.2 (Esteban et al., 2018), which is based on Nipype 1.8.5 (Gorgolewski et al., 2011).

##### Anatomical data

The T1-weighted (T1w) image was corrected for intensity non-uniformity (INU) with N4BiasFieldCorrection (Tustison et al., 2010), distributed with ANTs 2.3.3 (Avants et al., 2008), and used as T1w-reference throughout the workflow. The T1w-reference was then skull-stripped with a Nipype implementation of the antsBrainExtraction.sh workflow (from ANTs). Brain tissue segmentation of cerebrospinal fluid (CSF), white-matter (WM) and gray-matter (GM) was performed on the brain-extracted T1w using fast (FSL 6.0.5.1; Zhang et al., 2001). Brain surfaces were reconstructed using recon-all (FreeSurfer 7.2.0; Dale et al., 1999). The brain mask estimated previously was refined to reconcile ANTs-derived and FreeSurfer-derived segmentations. Volume-based spatial normalization to two standard spaces (MNI152NLin6Asym, MNI152NLin2009cAsym) was performed through nonlinear registration with antsRegistration (ANTs 2.3.3), using brain-extracted versions of both T1w reference and the T1w template.

##### Functional data

For each blood-oxygen level dependent (BOLD) run found per subject, the following preprocessing was performed. First, a reference volume and its skull-stripped version were generated for head motion correction. Head-motion parameters with respect to the BOLD reference (transformation matrices, and six corresponding rotation and translation parameters) were estimated before any spatiotemporal filtering using mcflirt (FSL 6.0.5.1; Jenkinson et al., 2002). The estimated fieldmap was then aligned with rigid-registration to the target EPI (echo-planar imaging) reference run and the field coefficients were mapped on to the reference EPI using the transform. BOLD runs were slice-time corrected to 0.346s (0.5 of slice acquisition range 0s-0.693s) using 3dTshift from AFNI (Cox & Hyde, 1997). The BOLD reference was then co-registered to the T1w reference using bbregister (FreeSurfer) which implements boundary-based registration (Greve & Fischl, 2009). This co-registration was configured with six degrees of freedom.

The BOLD time-series were resampled into standard space, generating a preprocessed BOLD run in MNI152NLin6Asym space. A reference volume and its skull-stripped version were generated using a custom methodology within fMRIPrep. The BOLD time-series were resampled onto the fsaverage surface (FreeSurfer). Grayordinates files (Glasser et al., 2013) containing 91k samples were also generated using the highest-resolution fsaverage as intermediate standardized surface space. All resamplings can be performed with a single interpolation step by composing all the pertinent transformations (i.e. head-motion transform matrices, susceptibility distortion correction when available, and co-registrations to anatomical and output spaces). Gridded (volumetric) resamplings were performed using antsApplyTransforms (ANTs), configured with Lanczos interpolation to minimize the smoothing effects of other kernels (Lanczos, 1964). Non-gridded (surface) resamplings were performed using mri_vol2surf (FreeSurfer).

#### Developmental sample postprocessing: (HCPD and HBN)

Preprocessed outputs were post-processed using the eXtensible Connectivity Pipelines–DCAN collaborative (XCP-D v0.3.0; Ciric et al., 2018; Mehta et al., 2024).

Specifically, before nuisance regression, the BOLD data were despiked, mean-centered, and linearly detrended. In total, 36 nuisance regressors were selected from the nuisance confound matrices of fMRIPrep output. These regressors included six motion parameters, global signal, mean white-matter signal, and mean cerebrospinal fluid (CSF) signal with their temporal derivatives, as well as the quadratic expansion of six motion parameters, tissues signals and their temporal derivatives (Ciric et al., 2017; Satterthwaite et al., 2013). Nuisance regressors were regressed from the BOLD data using linear regression implemented in Scikit-Learn 1.1.3 (Pedregosa et al., 2011). The residual time series were then band-pass filtered to retain signals within the 0.01–0.08 Hz frequency band. Finally, processed functional time series were extracted using Connectome Workbench (Glasser et al., 2013) for the Schaefer-400 atlas (Schaefer et al., 2018), yielding parcellated time series for 400 cortical regions. A lower-resolution Schaefer-200 parcellation was also used for sensitivity analyses.

#### Young adult sample (HCPYA)

All HCPYA fMRI data were preprocessed using the HCP Minimal Preprocessing Pipelines (Glasser et al., 2013), which include the same major steps as above: gradient distortion correction, field-map distortion correction, boundary-based registration, bias-field correction, and whole-brain intensity normalization.

The minimally preprocessed HCP outputs were then post-processed using XCP-D v0.9.1, following a workflow closely analogous to that used for the developmental sample. Framewise displacement was computed using the same formula (Power et al., 2014), and 36 nuisance regressors (“36P”) were selected following the same definitions as above (Ciric et al., 2017; Satterthwaite et al., 2013). The BOLD data were converted to NIfTI format, despiked with AFNI’s 3dDespike, and converted back to CIFTI format.

Denoising was performed using a regression approach based on Nilearn’s implementation. Time series and confounds were band-pass filtered (0.01–0.08 Hz) using a second-order Butterworth filter, and nuisance regression was applied to the filtered confounds. The denoised BOLD data were then used to extract parcellated time series for the Schaefer-400 atlas using Connectome Workbench. As with the developmental sample, a Schaefer-200 parcellation was also generated for sensitivity analyses.

### Intrinsic timescale estimated using the sum of positive autocorrelation

Fully processed parcellated fMRI time series were used to estimate intrinsic timescale for each brain region (i.e., parcel) and participant. Specifically, we first normalized the fMRI time series at each parcel using a z-score normalization; note that this standardization does not affect autocorrelation-based estimates. We then estimated an autocorrelation function (ACF) for the normalized time series using the Time Series Analysis (*tsa*) tool from *statsmodels* (Seabold & Perktold, 2010) in Python. Next, we identified the time point where ACF crosses zero autocorrelation value for the first time (i.e., first zero-crossing time point) and calculated the sum of all positive ACF values up to the first zero-crossing point. To obtain an estimate of the intrinsic timescale in seconds, we multiplied the sum of positive ACF values by the corresponding TR for each dataset (HCPD TR=800ms; HBN TR=800ms; HCPYA TR=720ms; Fulcher et al., 2013; Fulcher & Jones, 2017; Watanabe et al., 2019). Note that the fMRI acquisition TR corresponds to the time step between consecutive time points in ACF. This procedure is highly similar to calculating the area under the curve of the positive part of the ACF curve (**Figure 1**). We repeated this procedure for each parcel and individual, generating a parcellated map of intrinsic timescale for each participant.

We restricted our analyses to low-motion individuals (mean FD < 0.2 mm; see “Participants” for details), given that measures of intrinsic timescale are sensitive to in-scanner motion and physiological noise and are more reliable in higher-quality data with low motion (Goldberg et al., 2024). In addition, we included data quality (i.e., mean FD) as a covariate in subsequent models to account for any residual confounding effects of motion (see “Generalized Additive Model (GAM)” for details).

### Intrinsic timescale estimated using lag-1 temporal autocorrelation

As a complementary estimate of intrinsic timescale, we calculated lag-1 temporal autocorrelation (lag-1 TA) for each brain region and participant, following the approach used by Shinn et al. (2023) and Bero et al. (2026). Lag-1 TA quantifies the similarity between a time series and a copy of itself shifted by one time point. In the context of fMRI, this corresponds to the correlation between the BOLD signal at time point ***t*** and the BOLD signal at the immediately following time point, ***t***+1, within the same brain region. Higher lag-1 TA values therefore indicate slower decorrelation, or greater temporal persistence, across consecutive time points, whereas lower values indicate faster decorrelation from one time point to the next.

Fully processed parcellated fMRI time series were first z-score normalized within each parcel. We then estimated lag-1 TA using the temporal_autocorrelation(x) function from the “spatiotemporal” Python package (Shinn et al., 2023). This function computes TA-**Δ**_1_, defined as the first-order temporal autocorrelation between consecutive time points. For each participant, the input was a parcel-by-time matrix, and the output was a vector of lag-1 TA values across parcels. Unlike the ACF-sum timescale measure, lag-1 TA captures *only* the first nonzero lag of the ACF rather than the area under the positive portion of the ACF. Accordingly, lag-1 TA was not integrated across multiple lags and was not multiplied by TR; it is a dimensionless autocorrelation coefficient. We repeated subsequent analyses using these parcellated lag-1 TA maps to compare regional developmental patterns with those estimated using the original ACF-sum timescale metric.

### Short- and long-lag contributions to intrinsic timescale

To examine whether the ACF-sum measure of intrinsic timescale was differentially driven by short-versus longer-lag autocorrelation structure, we decomposed the positive ACF area into short- and long-lag components. First, using the normalized parcellated fMRI time series, we estimated the ACF for each parcel and participant as described above. We then computed a group-average ACF by averaging ACF curves across all parcels and participants. To identify a data-driven lag separating the steep early decay of the ACF from the slower positive tail, we fit a two-piece linear model to the positive portion of the group-average ACF. Specifically, for each candidate lag ***k***, we fit one linear function to the early-lag portion of the group-average ACF and a second linear function to the later-lag portion. We selected the lag ***k*** that minimized the total sum of squared errors across the two fitted line segments. Lag 0 was excluded from this elbow estimation because the ACF at lag 0 is fixed at 1. We searched candidate elbow lags from ***k*** = 3 to 47 within the positive group-average ACF, corresponding to 2.4-37.6 s in HCPD (TR = 0.8 s). The two-piece linear fit identified an elbow at lag ***k*** = 5, corresponding to 4.0 s in HCPD.

Using this single group-level elbow lag ***k***, we then returned to the participant- and parcel-level ACFs and calculated two complementary timescale estimates. The short-lag component, Int_short_, was calculated as the sum of ACF values from lag 0 through lag ***k***, bounded by the first zero-crossing point when the ACF crossed zero before lag ***k***. The long-lag component, Int_long_, was calculated as the sum of positive ACF values after lag ***k*** and up to the first zero-crossing point. Both components were multiplied by the dataset-specific TR to express them in seconds. This decomposition ensured that Int_short_ and Int_long_ summed to yield the original positive ACF-sum intrinsic timescale estimate for each parcel and participant. We used this approach to test whether short- and long-lag ACF components captured distinct aspects of development, and specifically whether lag-1 autocorrelation preferentially reflected the short-lag component of the ACF.

### Generalized Additive Model (GAM)

We used generalized additive models (GAMs) to assess linear and nonlinear associations with age in fMRI intrinsic timescale (Pomponio et al., 2020; Sydnor et al., 2023; Luo et al, 2024). The analysis was performed using the “mgcv” package in R 4.2.2. We included age as a smooth term and sex and data quality (i.e., in-scanner motion quantified by mean FD) as linear covariates in each GAM. We used Restricted Maximum Likelihood (REML) to estimate the smoothing parameters and set the maximum basis complexity to 3 for the smooth age term to avoid overfitting. Specifically, GAMs were formulated as ‘mgcv::gam(feature ∼ s(age, k=3, fx=F) + factor(sex) + data_quality)’. We performed two sets of analysis: (1) whole-brain GAMs, where we assessed developmental patterns of the whole-brain average intrinsic timescale; (2) regional GAMs, where we examined developmental patterns of regional intrinsic timescale.

The effect size of associations between intrinsic timescale and age were estimated as partial *R*^2^. Partial *R*^2^ was quantified as the difference in Sum of Squared Errors (SSE) between the full model that included the smooth age term and a reduced model with no age term (i.e., only including model covariates) normalized by SSE of the reduced model. Normalization by SSE of the reduced model highlights the relative contribution of the predictor (e.g., age) to the reduced model. We used a signed version of partial *R*^2^, such that the sign reflected the directionality of observed effects (increase or decrease in intrinsic timescale with increasing age). To obtain the directionality of partial *R*^2^, we calculated the mean derivative of the model fit for the smooth term (i.e., age). A positive sign was assigned to partial *R*^2^ if the mean derivative was positive, reflecting an overall increasing trend between the intrinsic timescale and age, whereas a negative sign was assigned to partial *R*^2^ for negative mean derivative. The statistical significance of the associations between intrinsic timescale and age was assessed using analysis of variance (ANOVA) between the full model and reduced model that excluded the smooth age term. Results were corrected for multiple comparisons by controlling for the false discovery rate (FDR correction; *Q*<0.05).

### Age of maximum alignment between age-related timescale differences and the S-A axis

To assess when age-related differences in intrinsic timescale were most strongly aligned with the cortical hierarchy, we estimated age-specific rates of timescale differences from the first derivative of each region’s GAM smooth function for age (Sydnor et al., 2023). These derivatives represent the estimated rate of timescale differences per one-year age interval. For each age from 8 to 22 years, we quantified the spatial association between regional derivative values and regional S-A axis ranks using Spearman’s rank correlation. To account for uncertainty in derivative estimates, we sampled 10,000 draws from the posterior distribution of the derivative for each regional GAM smooth and computed the age-specific correlation between derivative values and S-A axis ranks for each draw. The median correlation across draws was used to summarize the age-specific alignment between timescale differences and the S-A axis, and the 95% credible interval was calculated from the distribution of sampled correlations. We then identified the age of maximum alignment as the age at which the derivative-to-S-A axis correlation was maximal for each posterior draw. The median and 95% credible interval of these ages were used to summarize the timing of maximum alignment between age-related timescale differences and the S-A axis.

### Data acquisition site harmonization

To evaluate whether the findings were influenced by acquisition site, we harmonized intrinsic timescale estimates across data acquisition sites using CovBat-GAM (Johnson et al., 2007; Fortin et al., 2017; Fortin et al., 2018; Pomponio et al., 2020; Chen et al., 2022). Harmonization was performed in R 4.2.2 using the covfam function from the ComBatFamily package. This approach extends ComBat/CovBat harmonization to account for both mean and covariance differences across batches. In this analysis, the study site was treated as the batch variable. Age was included as a smooth term to preserve linear and nonlinear age-related effects, and sex was included as a linear covariate. To assess whether harmonization influenced our findings, we repeated the main analyses before and after site harmonization and compared the resulting whole-brain age associations, regional timescale maps, and regional age-effect maps.

### Sensorimotor-association timescale differentiation

To test whether timescale differentiation between sensorimotor and association cortex increased with age, we performed two supplementary analyses in HCPD. First, we conducted a within-subject distance-based analysis. Cortical regions were divided into lower-ranking sensorimotor and higher-ranking association regions based on their position along the S-A axis. For each participant, we computed pairwise absolute timescale differences among regions within the same end of the hierarchy and across sensorimotor-to-association regions. We then compared across-hierarchy differences to within-hierarchy differences using a two-sample *t*-test, yielding a subject-level *t*-value indexing sensorimotor-association timescale differentiation. We then tested whether this differentiation measure varied with age using a GAM with the same model structure as the main analysis.

Second, we performed a complementary analysis in which regional timescale values were directly compared between sensorimotor and association cortex for each participant using a two-sample *t*-test. The resulting subject-level *t*-values were then modeled as a function of age using the same GAM framework. For both analyses, effect size was quantified using partial *R*^2^ and significance was assessed by comparing the full model including age to a reduced model containing only covariates.

### Spatial null model

We used spatial autocorrelation-preserving null models (i.e., “spin” tests) to compare cortical patterns between different features or datasets (Alexander-Bloch et al., 2018; Markello et al., 2021). Specifically, we generated null brain maps with a randomized spatial distribution while preserving the spatial autocorrelation inherent to the data using the neuromaps toolbox (Markello et al., 2022). The randomization was implemented by applying 10,000 randomly sampled rotations to the spherical projections of the data for the Schaefer atlas (Alexander-Bloch et al., 2018; Markello et al., 2022). The rotations were applied to one hemisphere and then mirrored to the other hemisphere.

The resulting randomized (“spun”) brain maps were used for two types of significance testing. First, to evaluate the significance of associations between two cortical maps (e.g., the correlation between age effects and the S–A axis), we generated a null distribution of association statistics (e.g., Spearman’s rank correlation coefficient) by correlating one empirical map with 10,000 spun versions of the other map. The statistical significance of the observed association was determined by comparing the empirical statistic to this null distribution using a two-tailed test. Second, for analyses testing differences in mean values between two distributions (e.g., *t*-tests), spun brain maps were used to generate a null distribution of mean differences. The empirical mean difference was then compared against this null distribution to assess statistical significance (two-tailed test).

## Code and data availability

All code and accompanying guidelines used to conduct the reported analyses are available on GitHub (https://pennlinc.github.io/shafiei_timescale) and Zenodo (https://doi.org/10.5281/zenodo.22799416). Ready-to-plot data and scripts that are required to generate the main and supplementary figures are also provided in the same Zenodo record (https://doi.org/10.5281/zenodo.22799416). Raw data used in the present study were obtained from publicly available HCPYA (van Essen et al., 2013; https://www.humanconnectome.org/study/hcp-young-adult), HCPD (Somerville et al., 2018; https://www.humanconnectome.org/study/hcp-lifespan-development), and HBN (Alexander et al., 2017; https://fcon_1000.projects.nitrc.org/indi/cmi_healthy_brain_network/) datasets.

## Supplementary figures

**Figure S1.**
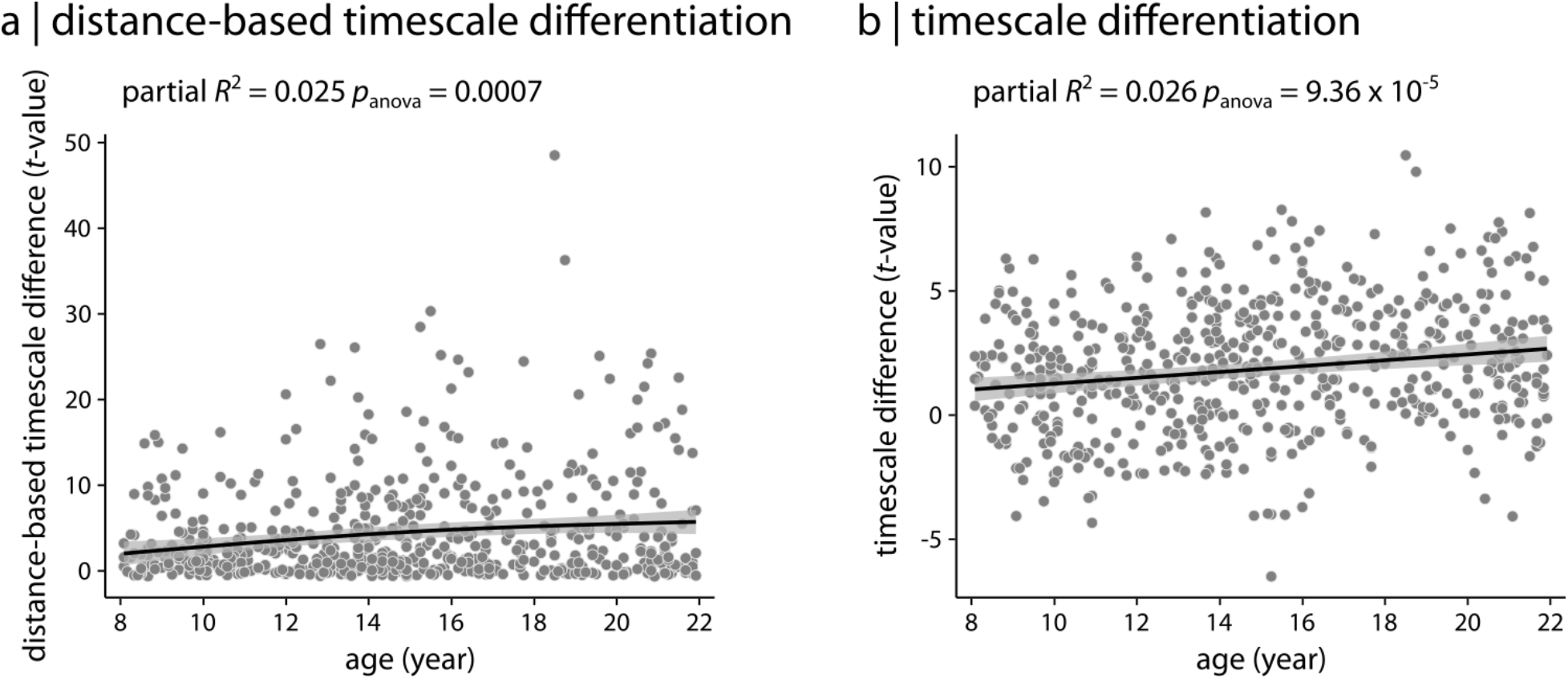
Sensorimotor-association timescale differentiation in HCPD. We tested whether timescale differentiation between sensorimotor and association regions increased with age. **(a)** In a within-subject distance-based analysis, cortical regions were divided into lower-ranking sensorimotor and higher-ranking association regions based on their position along the S-A axis. For each participant, pairwise timescale differences were computed within sensorimotor/association regions and across sensorimotor-to-association regions. Across-hierarchy differences were compared to within-hierarchy differences using a two-sample *t*-test, generating a subject-level estimate of timescale differentiation. This measure showed a significant positive association with age using a GAM (partial *R*^2^ = 0.025, *p*_anova_ = 0.0007). **(b)** In a complementary analysis, sensorimotor and association timescale values were directly compared within each participant using a two-sample *t*-test. The resulting subject-level *t*-values also showed a significant positive association with age using a GAM (partial *R*^2^ = 0.026, *p*_anova_ = 9.36 x 10^-5^). Each data point represents an individual participant. GAM fits are shown with black lines. Data and code needed to regenerate this figure can be found at https://doi.org/10.5281/zenodo.22799416.

**Figure S2.**
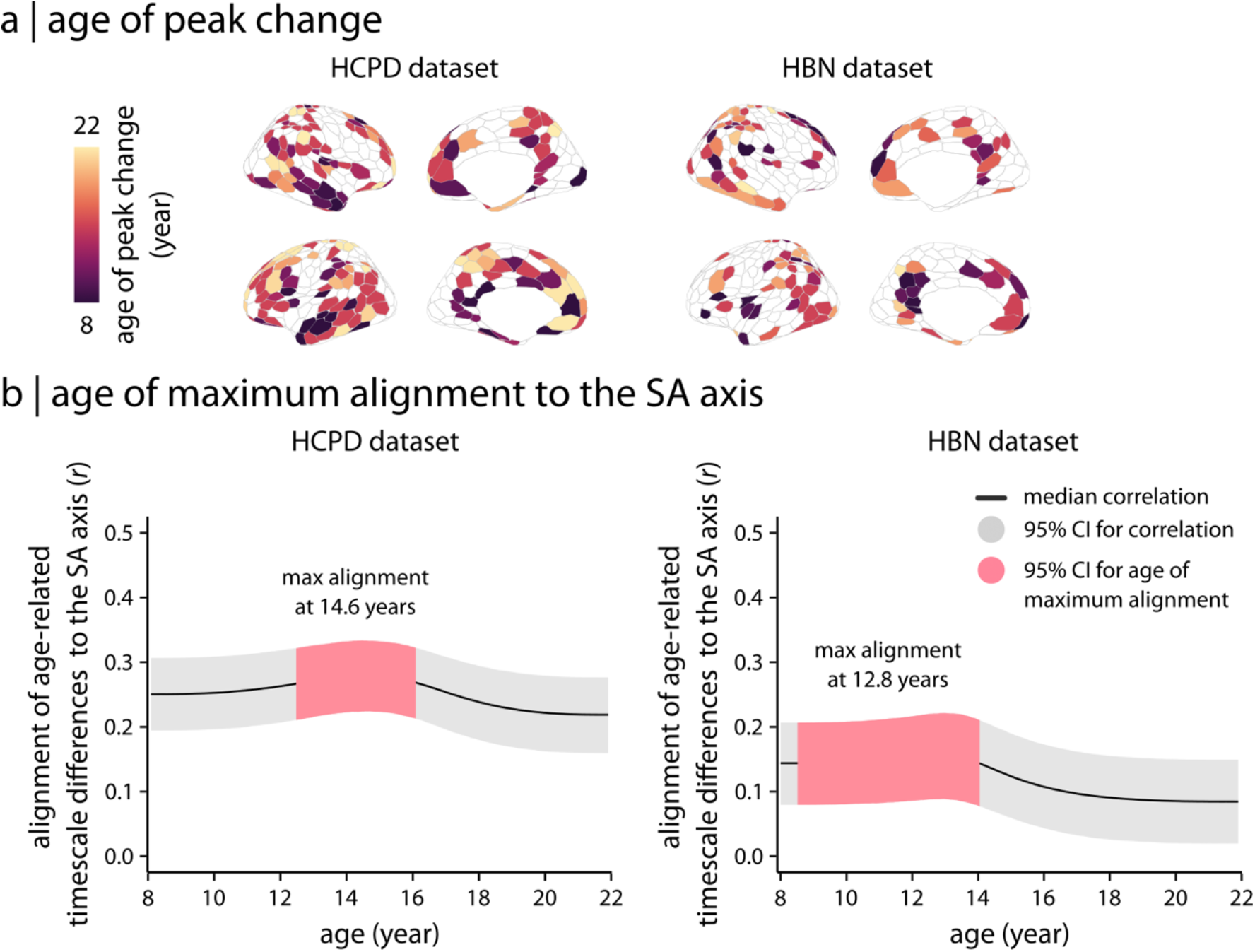
Timing of age-related differences in intrinsic timescale across two developmental cohorts. **(a)** Derivatives of the intrinsic timescale regional developmental patterns (Figure 3d and Figure 4d) were calculated as finite differences between consecutive time points to estimate the age at which timescale development demonstrated maximal change. Specifically, the age of peak change was estimated as the age where the derivative was greatest in absolute magnitude. **(b)** To estimate the age at which age-related timescale differences were maximally aligned with the S-A axis, we calculated the first derivative of each region’s GAM smooth function for age. These derivatives represent the estimated rate of timescale differences per one-year age interval. At each age, we quantified the spatial correlation between regional derivative values and regional S-A axis ranks. To account for uncertainty in derivative estimates, we sampled 10,000 draws from the posterior derivative distribution and calculated the derivative-to-SA axis correlation for each draw. The black line shows the median correlation across draws, and the gray band shows the 95% credible interval. The pink band shows the 95% credible interval for the age of maximum alignment. Maximal alignment occurred at 14.6 years in HCPD and 12.8 years in HBN. Data and code needed to regenerate this figure can be found at https://doi.org/10.5281/zenodo.22799416.

**Figure S3.**
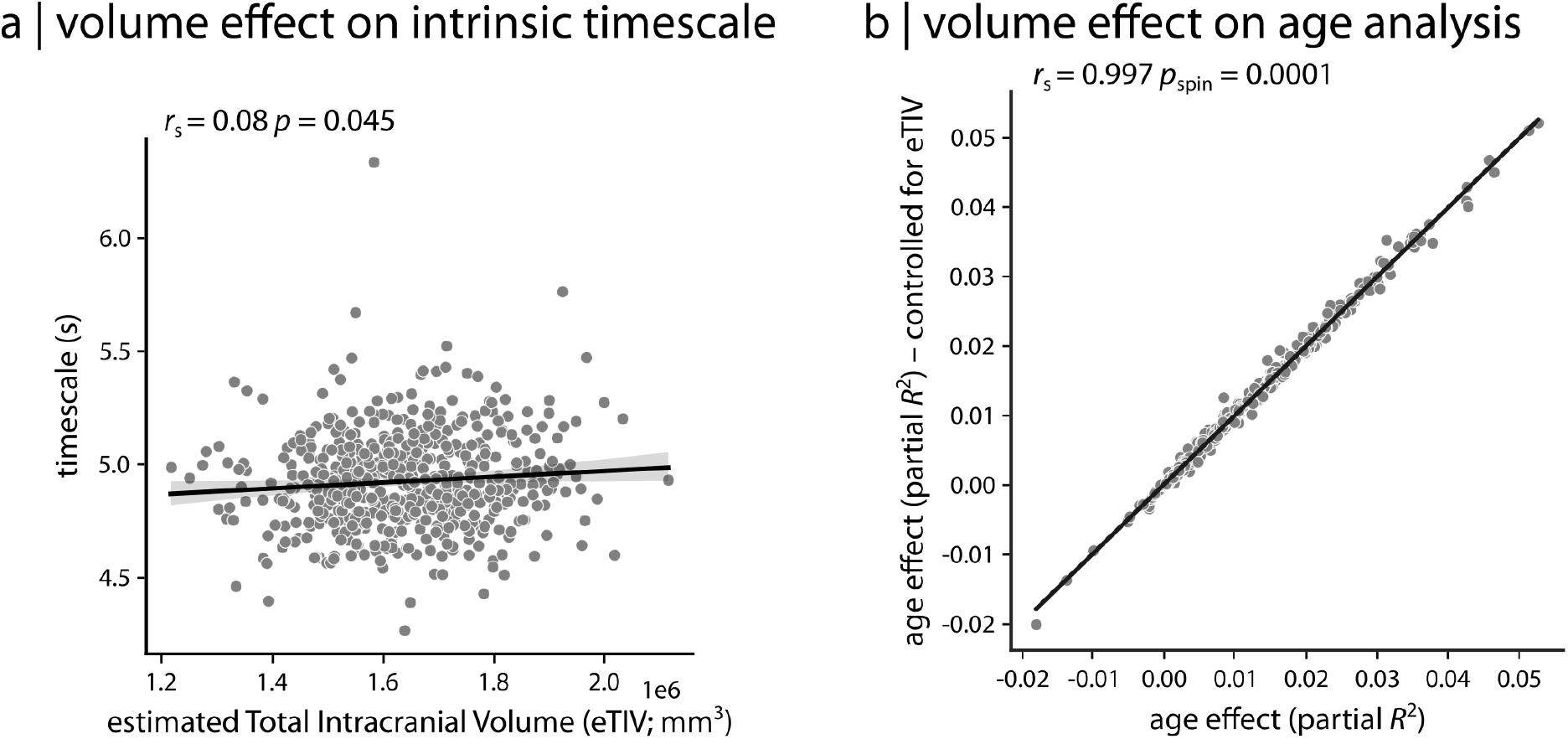
Effects of total brain volume on intrinsic timescale and its developmental patterns. **(a)** To ensure that findings were independent from changes in brain size during development, we directly examined the relationship between estimated Total Intracranial Volume (eTIV) and intrinsic timescale in the HCPD dataset and found a weak association between the two. Each data point in the scatter plot represents an individual participant. **(b)** To assess whether developmental patterns of intrinsic timescale were independent from changes in brain size, we repeated the age analysis (i.e., GAMs) while controlling for eTIV as a model covariate. The age effects (i.e., partial *R*^2^) were consistent with the original analysis. Each data point in the scatter plot represents a brain region from the Schaefer 400 atlas. *r*_s_ denotes Spearman’s rank correlation coefficient. Linear regression lines are added for visualization purposes only. Data and code needed to regenerate this figure can be found at https://doi.org/10.5281/zenodo.22799416.

**Figure S4.**
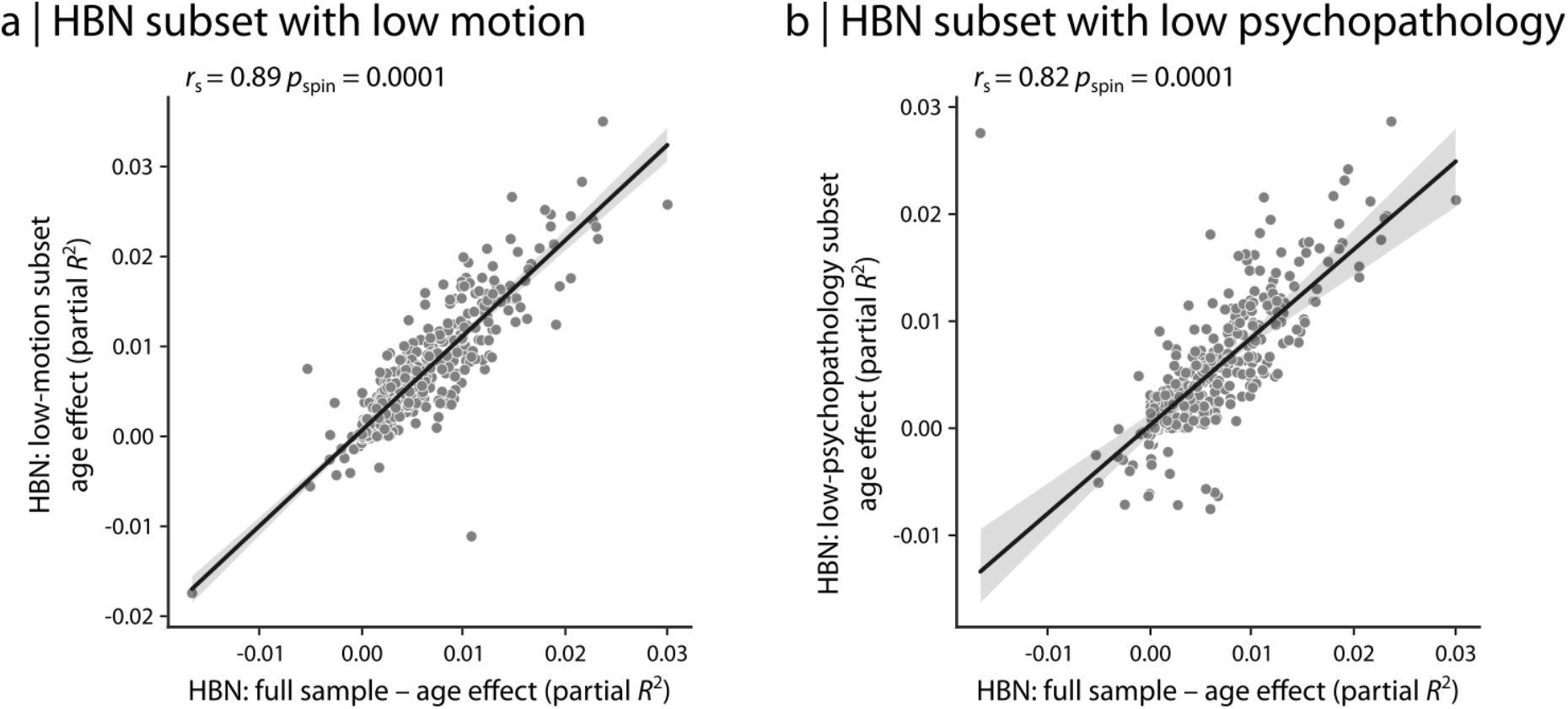
Consistent findings in two subsets of the HBN dataset. Given that HBN is a more heterogeneous sample compared to HCPD and includes help-seeking individuals with higher levels of psychopathology, we repeated all the analyses using two subsets of HBN individuals (each with *N*=600; this approximately matches the HCPD sample size) with **(a)** low motion and **(b)** low psychopathology (i.e., *p*-factor). In both cases, the developmental patterns in intrinsic timescale (i.e., partial *R*^2^) were consistent with the full HBN sample. Each data point in the scatter plot represents a brain region from the Schaefer 400 atlas. *r*_s_ denotes Spearman’s rank correlation coefficient. Linear regression lines are added for visualization purposes only. Data and code needed to regenerate this figure can be found at https://doi.org/10.5281/zenodo.22799416.

**Figure S5.**
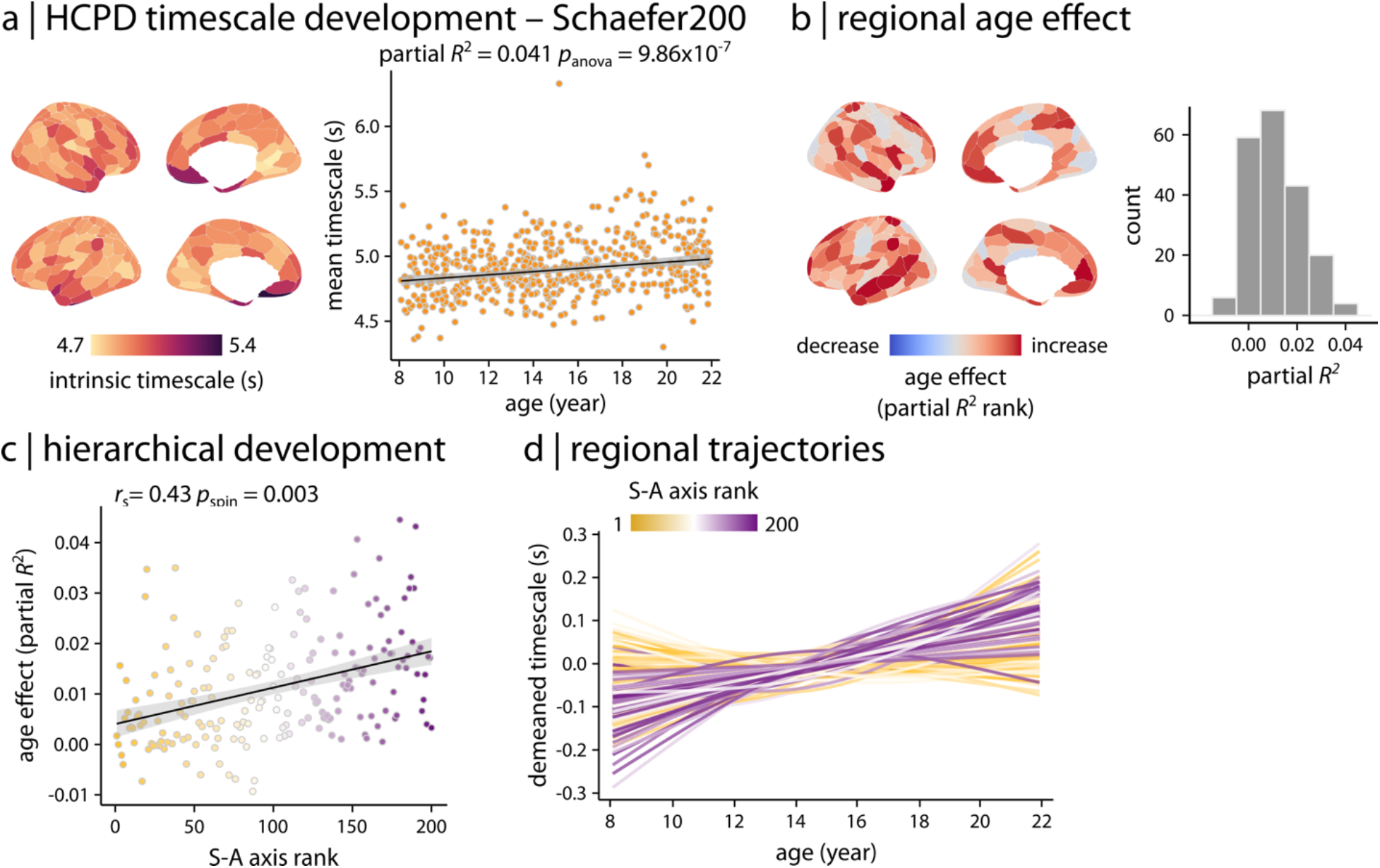
Consistent findings using the Schaefer 200 atlas. Developmental patterns of intrinsic timescale were replicated in the Schaefer 200 atlas in HCPD. **(a)** Consistent with the finding with the Schaefer 400 atlas (Figure 3), GAM results demonstrated that average timescale increases during development in youth (partial *R*^2^ = 0.041, *p*_anova_ = 9.86 x 10^-7^). **(b)** Region-wise GAMs identified heterogeneous age effects (i.e., partial *R*^2^) across the cortex. **(c)** Age effects on intrinsic timescale were hierarchically organized along the S–A axis. Significance of the association between age effects and S–A axis rank was assessed using 10,000 spin tests. *r*_s_ denotes Spearman’s rank correlation coefficient. Linear regression line is added for visualization purposes only. **(d)** Regional trajectories of developmental patterns in intrinsic timescale were obtained from region-wise GAM results (as shown in panel **b**). Each line corresponds to the model fit of each cortical region and is colored based on the region’s rank along the S–A axis. Similar to the findings with the Schaefer 400 atlas, the results demonstrate that intrinsic timescale in association regions increases during development in youth while it remains relatively stable in sensorimotor regions. Data and code needed to regenerate this figure can be found at https://doi.org/10.5281/zenodo.22799416.

**Figure S6.**
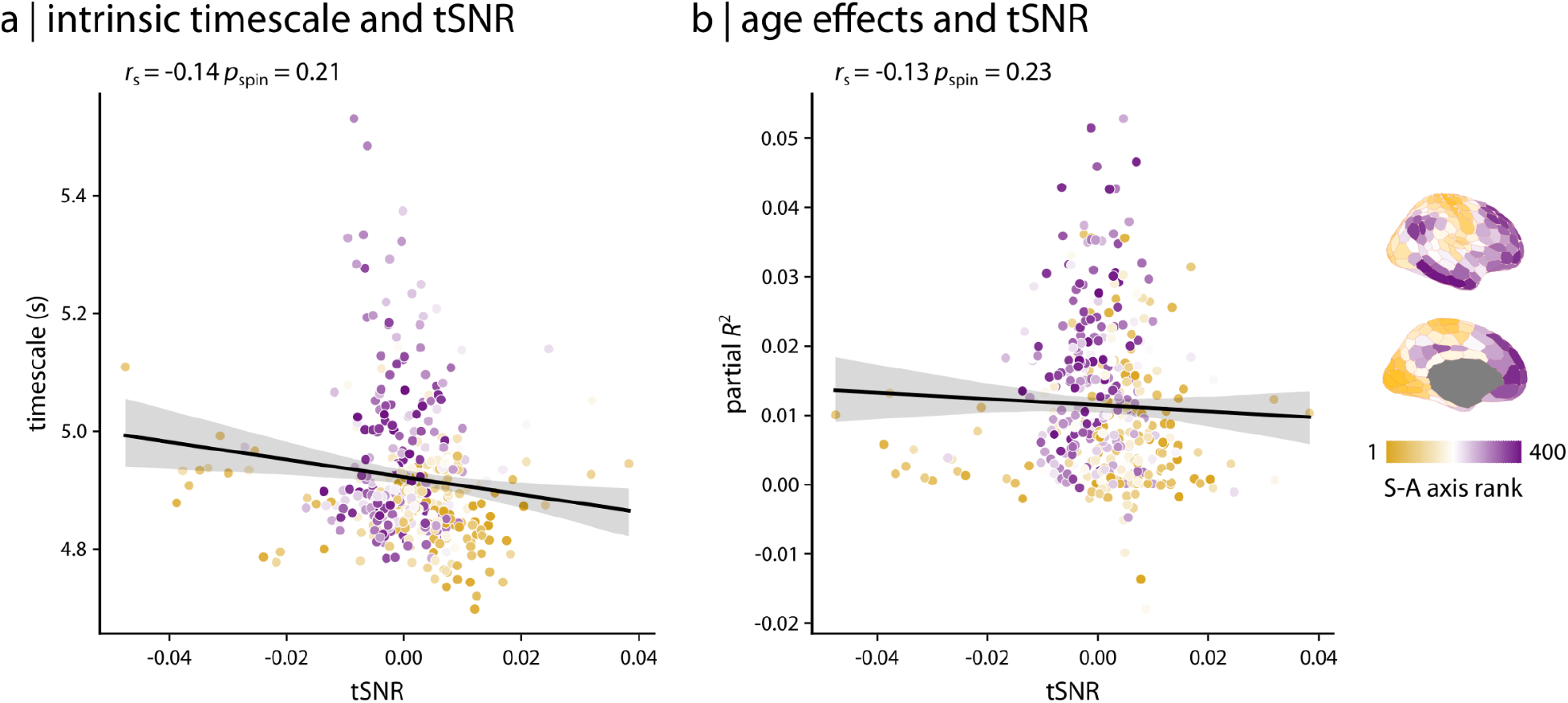
Findings were independent from temporal signal-to-noise ratio (tSNR) To test whether the findings were sensitive to signal-to-noise ratio (SNR) of the fMRI signal, we estimated temporal SNR (tSNR) as the ratio of the time-series mean to standard deviation for each region and participant in HCPD. Neither **(a)** the intrinsic timescale nor **(b)** the age effects (partial *R*^2^) were significantly associated with tSNR. Each data point in the scatter plot represents a brain region from the Schaefer 400 atlas. The data points are colored based on their rankings on the S–A axis. *r*_s_ denotes Spearman’s rank correlation coefficient. Linear regression lines are added for visualization purposes only. Data and code needed to regenerate this figure can be found at https://doi.org/10.5281/zenodo.22799416.

**Figure S7.**
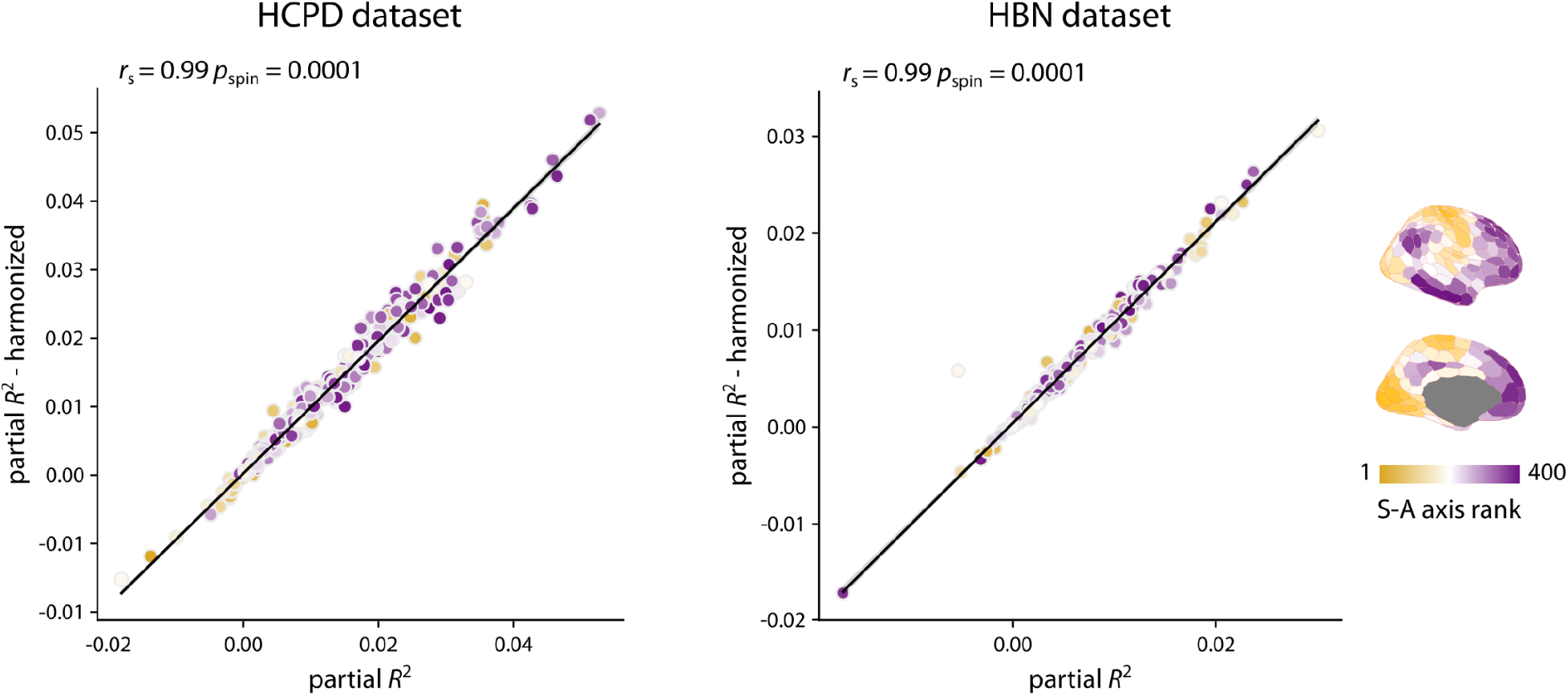
Findings were robust to site harmonization. To evaluate whether the findings were influenced by acquisition sites, we harmonized intrinsic timescale estimates across study sites using CovBat-GAM. Age effects (i.e., partial *R*^2^) were highly consistent before and after site harmonization. Each data point in the scatter plots represents a brain region from the Schaefer 400 atlas. The data points are colored based on their rankings on the S-A axis. *r*_s_ denotes Spearman’s rank correlation coefficient. Significance was assessed using 10,000 spin tests. Linear regression lines are added for visualization purposes only. Data and code needed to regenerate this figure can be found at https://doi.org/10.5281/zenodo.22799416.

**Figure S8.**
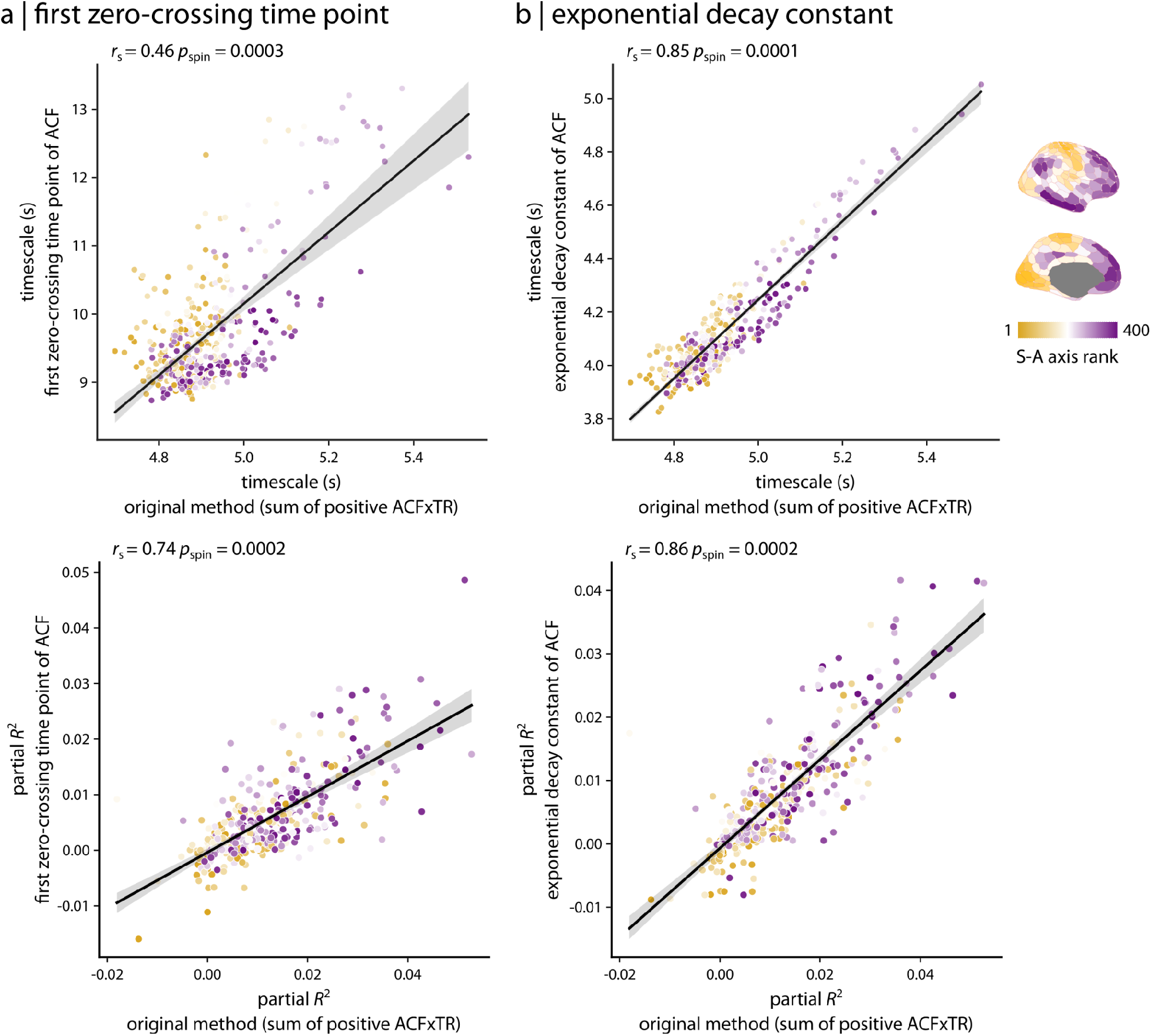
Convergent results with methodological sensitivity analysis in quantifying intrinsic timescale. To test whether intrinsic timescale estimates were sensitive to methodological choices, we quantified the intrinsic timescale in HCPD using two other approaches. **(a)** The intrinsic timescale was quantified as the first zero-crossing time point of the fMRI time series autocorrelation functions (ACF). **(b)** The intrinsic timescale was quantified as the exponential decay constant from an exponential fit to ACF. Both approaches generated intrinsic timescale maps consistent with the original method. Additionally, the developmental patterns were consistent with the original analysis. Each data point in the scatter plot represents a brain region from the Schaefer 400 atlas. The data points are colored based on their rankings on the S–A axis. *r*_s_ denotes Spearman’s rank correlation coefficient. Linear regression lines are added for visualization purposes only. Data and code needed to regenerate this figure can be found at https://doi.org/10.5281/zenodo.22799416.

**Figure S9.**
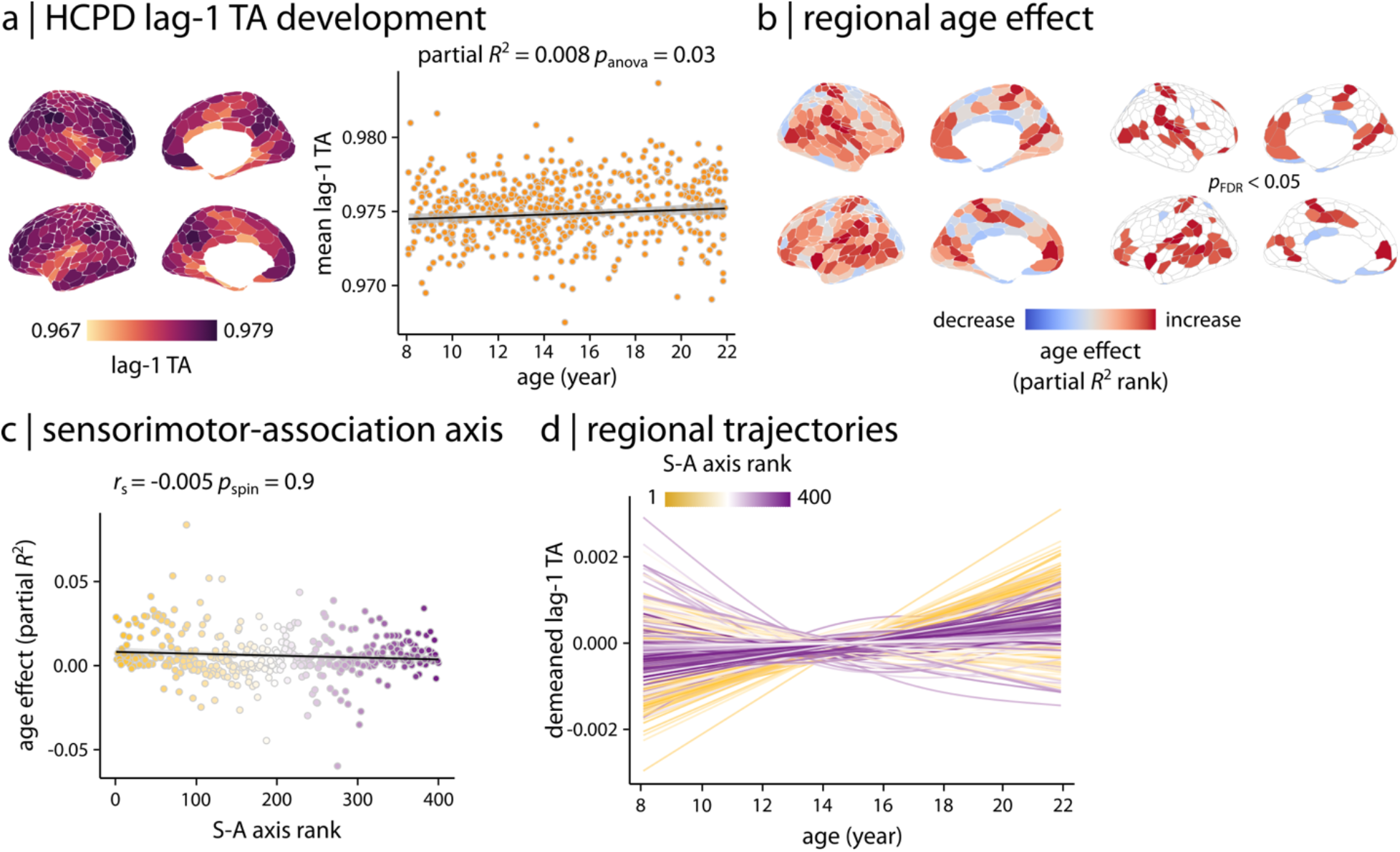
Developmental patterns of lag-1 temporal autocorrelation in HCPD. Lag-1 temporal autocorrelation (lag-1 TA) was estimated in HCPD as a complementary measure of intrinsic timescale. **(a)** Cortical map of lag-1 TA and relationship between whole-brain average lag-1 TA and age. Each data point in the scatter plot represents an individual participant (partial *R*^2^ = 0.008, *p*_anova_ = 0.03). **(b)** Region-wise GAMs identified heterogeneous age effects (i.e., partial *R*^2^) across cortex. FDR-corrected maps show regions with significant age effects (*p*_FDR_ < 0.05; 28.2% of regions with significant age effects). **(c)** Age effects on lag-1 TA were not significantly associated with the S-A axis. Significance of the association between age effects and S-A axis rank was assessed using 10,000 spin tests. *r*_s_ denotes Spearman’s rank correlation coefficient. Linear regression line is added for visualization purposes only. **(d)** Regional trajectories of developmental patterns of lag-1 TA were obtained from region-wise GAM results. Each line corresponds to the model fit of each cortical region and is colored based on the region’s rank along the S-A axis. Data and code needed to regenerate this figure can be found at https://doi.org/10.5281/zenodo.22799416.

**Figure S10.**
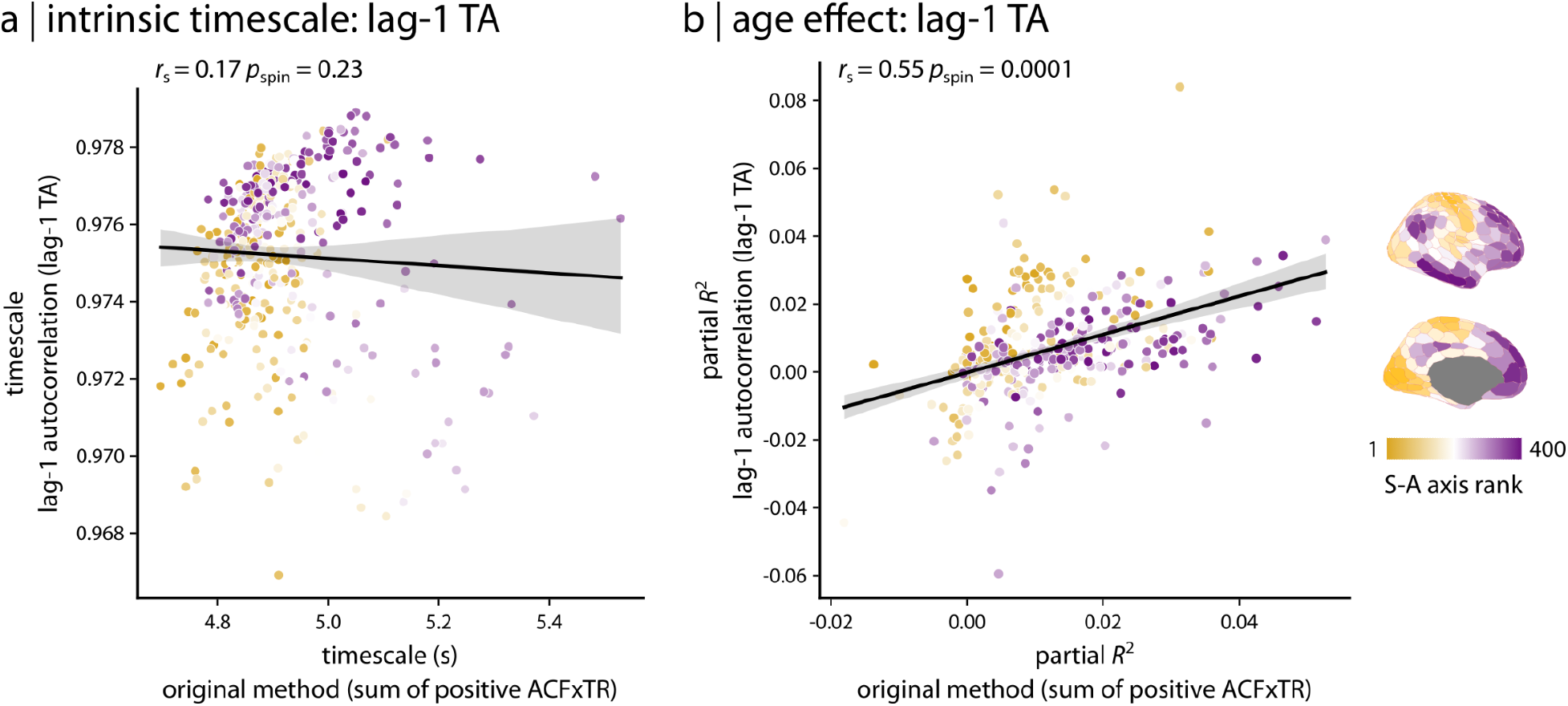
Relationship between ACF-sum intrinsic timescale and lag-1 temporal autocorrelation. We compared regional maps of intrinsic timescale estimated using the original ACF-sum method and lag-1 temporal autocorrelation (lag-1 TA) in HCPD. **(a)** Regional intrinsic timescale maps estimated using the two approaches were not significantly correlated. **(b)** In contrast, regional age effects (i.e., partial *R*^2^) estimated using the two approaches were significantly correlated. Each data point in the scatter plot represents a brain region from the Schaefer 400 atlas. The data points are colored based on their rankings on the S-A axis. *r*_s_ denotes Spearman’s rank correlation coefficient. Significance was assessed using 10,000 spin tests. Linear regression lines are added for visualization purposes only. Data and code needed to regenerate this figure can be found at https://doi.org/10.5281/zenodo.22799416.

**Figure S11.**
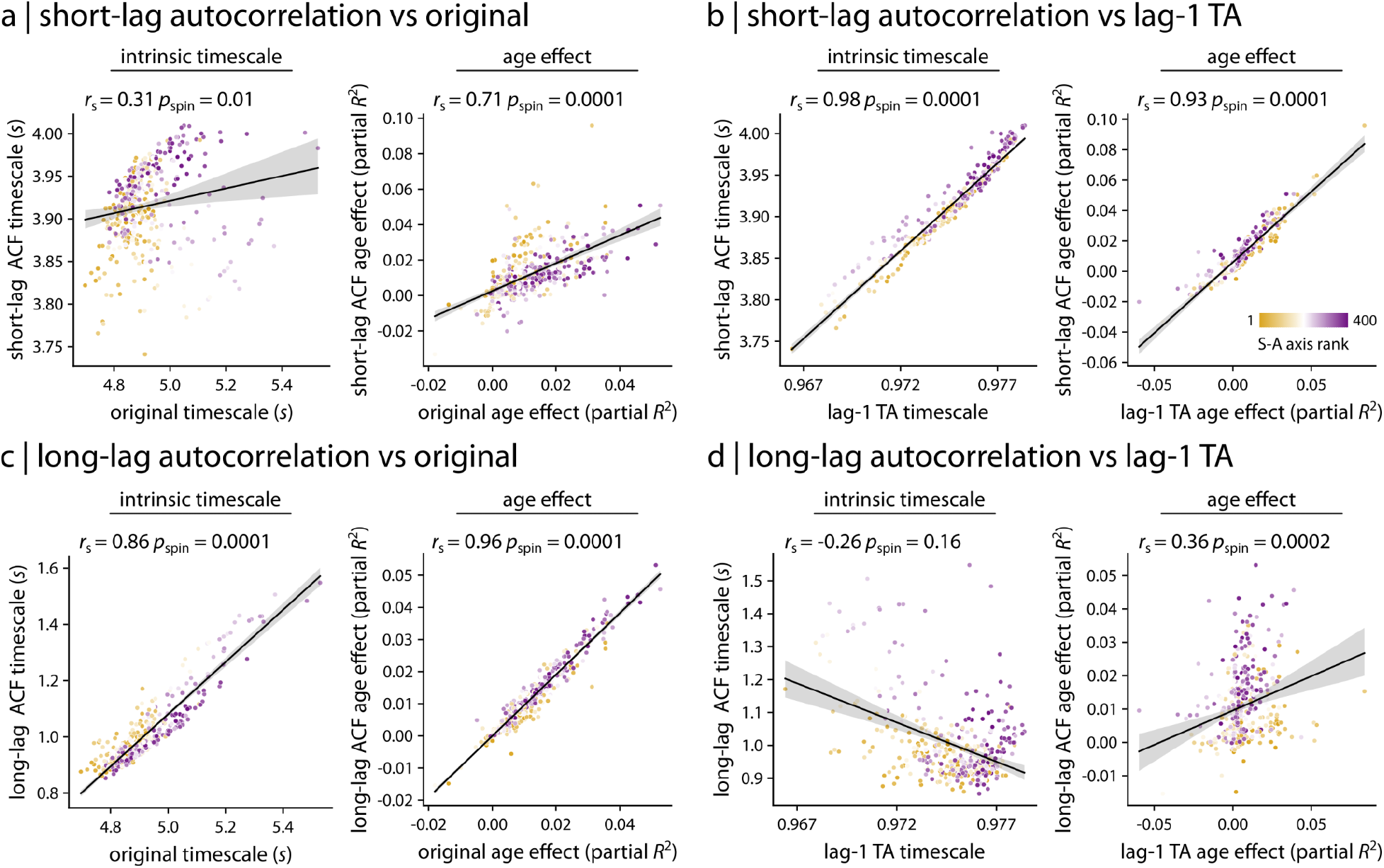
Short- and long-lag contributions to ACF-sum intrinsic timescale. To examine whether ACF-sum intrinsic timescale was differentially driven by short-versus longer-lag autocorrelation structure, we decomposed the positive ACF area into short-lag and long-lag components. **(a)** The short-lag component was modestly associated with the intrinsic timescale estimated using the original ACF-sum method. The regional age effects from the short-lag component were moderately associated with age effects from the original method. **(b)** In contrast, the short-lag component was highly associated with lag-1 temporal autocorrelation (lag-1 TA), both for regional timescale maps and regional age effects. **(c)** The long-lag component and its age effects were strongly associated with the original findings estimated using the ACF-sum method. **(d)** In contrast, the long-lag component was not significantly associated with lag-1 TA in regional timescale maps and was only weakly associated with lag-1 TA regional age effects. Each data point in the scatter plot represents a brain region from the Schaefer 400 atlas. The data points are colored based on their rankings on the S-A axis. *r*_s_ denotes Spearman’s rank correlation coefficient. Significance was assessed using 10,000 spin tests. Linear regression lines are added for visualization purposes only. Data and code needed to regenerate this figure can be found at https://doi.org/10.5281/zenodo.22799416.

